# Temperate phage microdiversity reflects infant gut microbiome maturation independent of chronic undernutrition

**DOI:** 10.64898/2026.05.06.723284

**Authors:** Laura Carolina Camelo Valera, Alejandro Reyes Muñoz, Corinne F. Maurice

## Abstract

The assembly and maturation of the infant gut microbiome is a critical developmental process. Yet the dynamics of the viral community, particularly in the context of stunting (chronic malnutrition) remain underexplored. Leveraging longitudinal fecal metagenomes from Zimbabwean infants with normal and stunted growth trajectories, we characterized the development of the gut bacterial and temperate phage communities from birth to 18 moths old. We found that infant gut temperate phages target hallmark early-life bacterial taxa, such as *Bifidobacteriaceae,* and exhibit an age-dependent maturation that parallels bacterial succession. Notably, both bacterial and temperate phage alpha diversity increased with age. This contrasts with previous studies focused on the extracellular viral fraction and highlights a strong coupling between prophage early-life dynamics and during bacterial gut colonization. Using abundance-based maturation models, we identified successional phases of colonization for both bacteria and their associated temperate viral clusters. Importantly, a viral microdiversity maturation model provided a stronger prediction of chronological age than viral abundance-based model, revealing within-phage genomic variation as a key signal of virome assembly, particularly around weaning. Contrary to findings in wasting (or severe acute malnutrition), stunted growth trajectories were not associated with a significant delay in either bacterial or temperate phage maturation. These results demonstrate that viral genomic variation is a new, informative dimension of early-life gut microbial assembly and that stunting may not impair infants’ gut maturation process.

**Importance:** The early-life period represents a critical window for the establishment of the gut microbial communities, a process that is often affected by environmental factors such as diet. While severe acute malnutrition (SAM) is known to delay bacterial maturation, the impact of chronic, moderate undernutrition, such as stunting is poorly understood. Stunting is a highly prevalent global health condition with irreversible consequences on long-term host health, yet its implications on gut microbiome assembly remain unclear. Our study provides novel insights into the maturation of temperate phages, which to prime the infant gut by colonizing alongside their bacterial hosts and acting as drivers of bacterial evolution via lysogeny. By demonstrating that viral strain-level (genomic) variation captures a stronger age-related signal than viral abundance, we identified an underexplored dimension of microbial assembly. The finding that stunting, in contrast to SAM, does not impact microbial maturation provides essential context for public health interventions and future studies addressing this condition.

## Introduction

The assembly of gut microbial communities begins at birth and, for the first three years of life, goes through a maturation process driven by the progressive arrival and establishment of new microorganisms from the mother and other environmental and dietary exposures^1,2^. Typically, the gut bacteriome follows a reproducible sequence matching infant developmental milestones: newborns are enriched in facultative anaerobes and human milk oligosaccharide (HMO) consumers such as *Bifidobacterium* and *Lactobacillus*^1,3,4^. These bacteria produce short-chain fatty acids (SCFA), critical for infant gut epithelial cells and immune system training^5,6^. With the introduction of solid foods, there is a bacterial succession and overall increase in gut bacterial richness, characterized by the expansion of fiber-degrading taxa such as *Faecalibacterium prausnitzii*, capable of metabolizing more complex dietary compounds^2,7^. In parallel, the infant gut virome, dominated by bacteriophages, also matures, with age-discriminatory phages identified and supporting a viral succession taking place^8–10^. Recent studies indicate that temperate phages are highly abundant and commonly induced during the first months of life from the first bacterial colonizers^11,12^. In contrast to bacteria, viral richness decreases, suggesting distinct maturation dynamics between these communities^13^.

The coordinated maturation of the gut bacteriome and virome is shaped by early-life diet and nutritional context, and in settings where undernutrition is prevalent, gut microbiome development follows altered trajectories^14,15^. For example, severe acute malnutrition (SAM), defined by a weight-for-height Z-score (WAZ) below -3, is associated with persistent delays in bacterial maturation^15^. These alterations extend to the gut virome: some Caudoviricetes phages have been identified as biomarkers of nutritional status, discriminating healthy infants from those with kwashiorkor or marasmus^9^. While SAM is a life-threatening form of undernutrition, chronic malnutrition remains the most prevalent form of undernutrition worldwide, particularly in Middle and Eastern Africa and Southern Asia^16^. Chronic malnutrition impacts approximately one in five children globally and results in impaired linear growth, or stunting, defined by a length-for-age Z-score (LAZ) less than -2 standard deviations from the median of the WHO child growth standards^16^. Stunting has received comparatively less mechanistic investigation than SAM, despite its high prevalence and irreversible consequences on physical growth, neurocognitive development, and long-term metabolic health^8,14,17^. Identifying whether there are gut microbial maturation patterns unique to stunting is therefore important.

Phages shape bacterial communities at multiple levels: through predation, they influence abundance and diversity, and through lysogeny, they drive bacterial evolution through horizontal gene transfer, reshaping the genetic pool of the bacterial community^18,19^. Foundational infant virome studies largely focused on the extracellular virome using virus-like particle (VLP) enrichment^9,10^. However, VLP-based and bulk metagenomics approaches capture distinct viral fractions, with only an estimated 8.5% overlap, reflecting the high prevalence of integrated prophages in infant gut microbiomes^13,20^. Consequently, bulk metagenomes provide complementary insights to VLP-based approaches, enabling high-resolution characterization of phage-host interactions and prophage dynamics during early-life development.

Across ecosystems, these interactions often operate at fine genomic resolution, with environmental conditions shaping whether phage pressure drives bacterial strain-level turnover or broader community change^21^. For example, spatial structuring and resource limitation tend to favor localized interactions and fluctuating selection dynamics, leading to locally adapted phage-bacteria strain pairs rather than repeated species-level shifts. In contrast, high-nutrient conditions may favor lysogenic strategies, which often exert weaker immediate effects on community composition, and are consistent with the “piggyback the winner” model^18,21,22^. Taken together, these patterns suggest that phages can also act as regulators of microbial microdiversity, rather than only shaping species-level composition. Applying this framework to infant gut development, we expect that as microbial maturation progresses and diets change, viral microdiversity increases, expanding phage functional potential (e.g., host range) and shaping co-evolutionary dynamics (arms-race versus fluctuating selection), and that chronic undernutrition disrupts these trajectories.

Viral microdiversity remains underexplored in early life, particularly in relation to nutritional status, despite its potential to reveal how within-species viral variation influences bacterial diversity. Here, we analyze a publicly available dataset of fecal bulk metagenomes from healthy and stunted infants in rural Zimbabwe, a region with a high burden of chronic undernutrition^14^. Because the original study focused on the effects of HIV exposure on bacteriome maturation, we isolated the role of nutritional status by restricting our analysis to infants born to HIV-negative mothers. Leveraging the prophage information contained in bulk metagenomes, we characterized prophage dynamics and examined viral maturation at the macro- and micro-diversity scales to evaluate if stunting is associated with altered bacterial or viral abundance and viral microdiversity trajectories and thus, by extension, with changes in phage-bacteria interactions that may influence the development of a healthy gut ecosystem^9,10,13^.

## Methods

### Data acquisition, preprocessing, and cohort description

We used a publicly available dataset originating from the Sanitation, Hygiene, Infant Nutrition Efficacy (SHINE; NCT01824940) trial^23^, which enrolled infants from two rural districts in Zimbabwe (Chirumanzu and Shurugwi) as part of improved Water, Sanitation and Hygiene (WASH) and Infant and Young Child Feeding (IYCF) interventions. Raw fecal metagenome sequencing data and corresponding cohort metadata from this study are deposited in the European Nucleotide Archive (ENA) under accession number PRJEB51728, which contains bulk fecal metagenomes from participating infants. The cohort metadata, including all epidemiologic data was retrieved from the second version of a public repository under the ZenodoID 7474876, from the R data structure (metaphlan.rds). The metadata included information on infant age, sex, WHO Z-scores for height/length-for-age (LAZ), weight-for-age (WAZ), and weight-for-height/length (WHZ), along with Middle Upper Arm Circumference (MUAC) and HIV status of the mother during pregnancy. The metadata was filtered to remove duplicate entries based on the sample identifier. The pregnancy HIV status variable (preg_hiv_status) was used to filter the samples, retaining only samples where the status was explicitly “Negative”. This step resulted in a subset of 484 samples from 236 infants never exposed to HIV (**Figure 1a**).

**Figure 1.**
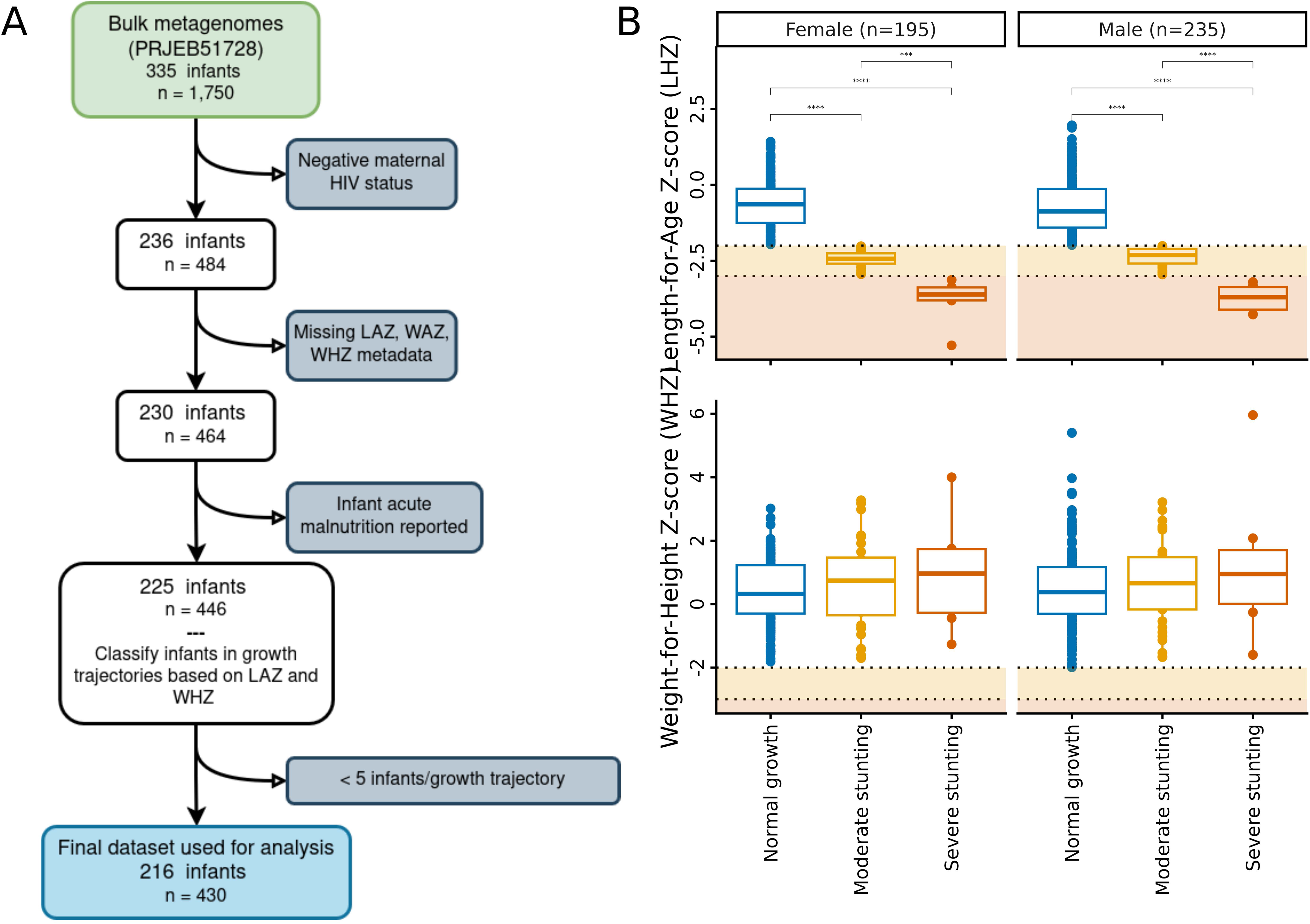
Sample selection workflow and anthropometric characterization. **(A)** Schematic of the decision-making process for sample selection. The SHINE^14,23^ cohort used for subsequent analyses initially comprised 1750 samples from 335 infants. Further data cleaning based on metadata and the removal of HIV-positive cases resulted in a final set of 430 samples from 216 infants later categorized in different growth trajectories. (B) Anthropometric measurements of the included set of infants. Wilcoxon signed-rank tests were used to compare LAZ and WHZ scores across nutritional statuses; only significant comparisons (∗p<0.05,∗∗p<0.01,∗∗∗p<0.001) are shown.

### Classification of growth status and longitudinal trajectories

Stool samples were collected from children in this cohort from birth to 600 days, with an average of approximately 1.96±0.97 samples per infant. Next, we stratified each sample into three growth groups based on reported LAZ on visit following WHO standards^16^: Normal growth (LAZ> -2; n = 356), moderate stunting (-3<= LAZ<=-2; n = 61), and severe stunting (LAZ < -3; n = 13). We further grouped samples at the infant level, individuals were classified into longitudinal trajectories based on their growth status across all visits. Trajectory groups represented by less than 5 infants were excluded for downstream analyses (10 groups comprising 16 samples). The remaining trajectories were consistent normal growth (n = 166 samples), decline to moderate stunting (n = 20), consistent moderate stunting (n = 18), recovery to normal growth (n = 6), consistent severe stunting (n = 9) **(Supplementary Fig 1)**. This resulted in a final subset of 430 samples from 219 infants.

### Bioinformatics pipeline for microbial communities characterization

#### Metagenomic data preprocessing

The bioinformatics pipeline to characterize gut bacterial and phage communities started with a read quality control and decontamination stage to ensure high-quality sequencing data. Raw sequencing reads were first processed for quality control (SLIDINGWINDOW:4:20 MINLEN:75 HEADCROP:10) and adapter removal using Trimmomatic^24^ (v0.39). Trimmed reads were subsequently mapped and filtered against the human reference genome (Homo_sapiens.GRCh38; GCA_000001405.29) to remove host contamination using Bowtie2^25^ (v2.4.2). The set of trimmed, human-decontaminated reads was subsequently used both for read-based bacterial taxonomic profiling and for viral identification following contig assembly.

#### Bacterial taxonomic profiling

To characterize the bacterial community composition, we used MetaPhlAn^26^ (v4.1.1). Human decontaminated reads were mapped against the MetaPhlAn^26^ database of genome markers from reference species genome bins: metaphlan4_db_v_Oct22. Alignments were performed using Bowtie2^25^ (v2.5.4) with MetaPhlAn’s default parameters. We utilized the -t rel_ab_w_read_stats flag to estimate relative abundances capturing read count statistics for each identified taxon. The resulting taxonomic profiles and abundance tables were used to determine the bacterial richness and community structure across all samples.

#### Viral contig assembly, identification and dereplication

For gut viral communities characterization, we assembled our clean reads set, then performed viral identification from the assembled contigs, dereplication, and abundance quantification. Human decontaminated reads were assembled for each sample individually using the metaSPAdes^27^ assembler (v3.15.5). Assembly was performed with 12 threads and a memory limit of 98 GB. Contigs from each sample’s assembly were screened for viral sequences using geNomad^28^ (v1.7.4). geNomad^28^ was used to classify viral and proviral sequences, which were then extracted into FASTA files. All identified viral contigs from the cohort were pooled, and preliminary dereplication was performed using BLASTn^29^ (v2.14.0) to calculate all-vs-all pairwise alignment statistics. Viral sequences were clustered at 95% Average Nucleotide Identity (ANI) across 85% target coverage using the anicalc.py and aniclust.py scripts from the CheckV^30^ suite (v1.0.1), to define the initial set of viral Operational Taxonomic Units (vOTUs). The resulting vOTU dataset was concatenated with the publicly available Gut Viral Database (GVD)^13^, downloaded from iVirus^31^. This combined contigs dataset was then subjected to a minimum sequence length filtering of 5kb and a second round of dereplication using the same 95% ANI and 85% coverage thresholds^32^. This step yielded a final, non-redundant reference set of 55,981 vOTUs.

#### vOTU read-based abundance quantification

The cleaned, human-decontaminated reads from all samples were mapped back to the final dereplicated vOTU reference set (study dataset + GVD) using Bowtie2^25^ (v2.4.2). This was performed using standard, non-competitive mapping. The resulting SAM files were converted and sorted into BAM format using SAMtools^33^ (v1.18). The coverage depth for each vOTU in every sample was calculated using SAMtools^33^ coverage. Prior to downstream analysis, the initial coverage file was pre-filtered to retain only vOTUs that met high-confidence criteria: coverage (breadth) greater than 75% and mean depth greater than 1 read per base^34^. Following these filtering steps, a final subset of 23,679 vOTUs remained present across all the samples and was used for all subsequent diversity and longitudinal modeling. In R^35^ (v4.4.0), the relative abundance of each vOTU within a sample was calculated by total-sum scaling as previously described^34^.

#### vOTUs characterization: quality check, host and replication cycle predictions

The final dereplicated vOTUs were classified into quality categories: Complete, High-quality, Medium-quality, Low-quality, and Not-determined, using CheckV^30^ (v1.0.1). The predicted replication cycles (i.e., virulent or temperate) of the vOTUs were determined using BACPHLIP^36^ (v0.9.6). A vOTU was classified based on the highest probability determined: virulent if virulent_prob>0.5; temperate if temperate_prob>0.5; no confidence if neither probability exceeded 0.5. To improve confidence in the final classifications, the replication cycle assignments were filtered by CheckV^30^ quality: only vOTUs classified as Complete, High-quality, or Medium-quality were retained. Virulent vOTUs that were not classified as complete were re-classified as “Non-temperate” to reflect their partial completeness, and vOTUs with “No Confidence” were excluded.

vOTUs host prediction was performed using iPHoP^37^ (v1.0.0). For vOTUs with multiple assignments, the entry with the highest confidence score was selected^38^. Host taxonomy was parsed to obtain classifications at the family, genus, and species level. The resulting host assignments were integrated with the replication cycle predictions. As we decided to focus only on temperate phage communities, we decided to filter out all the non-temperate and virulent vOTUs from subsequent analyses.

#### Viral taxonomic clustering

The final dereplicated set of vOTUs was taxonomically clustered based on shared protein content using vConTACT3^39^ (v3.1.6) with default settings. To facilitate downstream analyses, vConTACT3 output (final_assignments.csv) was processed using the phu^40^ utility simplify-taxa with the –add-lineage flag to generate Viral Clusters (VCs) nomenclature. This step standardized long taxonomic strings into compact, hierarchical lineage codes while maintaining consistency with the International Committee on Taxonomy of Viruses (ICTV) nomenclature^41^ (i.e.Caudoviricetes:NO3:NF2:NG1). Taxonomic assignments were resolved to the deepest possible rank, for vOTUs where higher-resolution classification (i.e. Genus) was unavailable, the next deepest possible taxonomic rank was retained (i.e. Family). For vOTUs lacking a taxonomic assignment (NA), the cluster was designated as a ’singleton’ followed by the unique vOTU identifier. Following this taxonomic annotation, vOTUs read-based abundances (see above) were aggregated at the VC level by summing the relative abundances of all vOTUs sharing the same lineage. The resulting matrix was merged with sample metadata (subjectID, age, stunted_status, etc.). All data manipulation and matrix construction downstream of the phu utility were performed in R^42^ (v4.5.1).

#### Viral nucleotide diversity calculations

We quantified within-sample nucleotide diversity using inStrain^43^ profile workflow (v1.8.0) on the BAM files obtained from the standard, non-competitive mapping of sample reads to the final dereplicated vOTU reference set. For each sample, we profiled coverage, SNVs, and gene-level diversity using --min_mapq 2, then we applied an additional quality filter for diversity estimates whereby viral contigs were only considered if they achieved ≥50% genome coverage (breadth) in such sample. Nucleotide diversity metrics were turned into a matrix with samples as rows and vOTUs as columns (absences set to zero), and were subsequently used as inputs for feature selection and nucleotide diversity-based maturation model development.

### Bioinformatic pipeline for microbial ecology and statistical modeling

#### Alpha diversity modeling

To model the longitudinal alpha diversity maturation of bacterial and temperate phage communities across different nutritional trajectories, we first assessed whether the sequencing depth correlated significantly with infant age. Initial linear modeling using the lm function in R stats package^42^ (v4.5.1) confirmed that both, total metagenomic read counts and the number of reads mapped to vOTUs, increased significantly with infant age across all nutritional status (p<0.05). Thus we included sequencing depth as a fixed factor when modeling alpha diversity for each community.

Several alpha diversity metrics were calculated using the vegan^44^ package in R (v2.7-3): Observed Richness (total number of distinct bacterial species or vOTUs identified per sample), Pielou evenness to assess how equally bacterial species or temperate vOTU abundances were distributed in each sample, and Shannon index. For the viral portion, we only considered vOTUs predicted as temperate. Now, to longitudinally model each of these metrics in each community, accounting for non-linear trends, we employed Generalized Additive Models (GAMs) using the mgcv^45^ package in R (v1.9-4). Separate models were applied for each alpha diversity metric in each community using the following global formula:

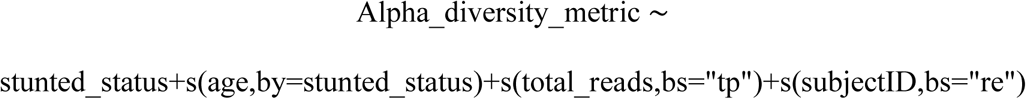

With this, age was modeled using a smooth term with thin plate regression splines, stratified by nutritional status to allow for unique maturation trajectories within each group. To account for variation in sequencing depth, total_reads was included as a smooth covariate. The longitudinal nature of the data and repeated measures within infants were accounted for by including subjectID as random effect smooth. All models were fitted using Restricted Maximum Likelihood (REML) to ensure unbiased parameter estimation.

To interpret and compare the models, Estimated Marginal Means (EMMs) were extracted using the emmeans^46^ package in R (v2.0.2). We evaluated the predicted diversity metrics across a 10-step age range (0 to 600 days), holding the sequencing depth constant at the mean value. Maturation trajectories were visualized by plotting the EMMs (Z-scores) against age. 95% confidence intervals were represented as ribbons or error bars to identify significant divergence between the normal growth reference group and transitioning or consistently stunted groups.

#### Data cleaning and feature engineering for normal growth maturation model

To define normal growth maturation models, we used the microbiome age^15^, defined as the age predicted based on microbial communities features, quantified either by relative abundance or nucleotide diversity. Using bacterial species and viral matrices from infants with consistent normal growth we built maturation models separately for each data modality (abundance- and nucleotide diversity–based) and for each organism (bacteria or temperate phages). We followed the steps outlined below:

##### Step 1. Pre-processing

To ensure data quality and biological relevance, we filtered out taxa with zero abundance across all samples and samples with a total mean depth of less than 1,000. For nucleotide diversity matrices, vOTUs were additionally filtered to retain only those with at least 50% genome coverage within a given sample. Each matrix was then transposed to have samples as rows (instances) and taxa as columns (predictors), taxa not present in a sample would appear as missing values, which were converted to zero, indicating an absence of that feature in the sample.

##### Step 2. Feature selection for maturation model

To characterize VCs and bacterial species that are associated with infant chronological age, and consequently gut microbiome maturation, we performed a feature selection step applying the Boruta^47^ algorithm (v9.0.0) built on Random Forest (500 trees) in each iteration (for a total of 500 iterations). This allowed us to reduce the model complexity by only using taxa that act like markers of maturation, so we model our healthy maturation reference curves using only taxa categorized as “Confirmed” or “Tentative” based on their Mean Decrease Gini importance (output from Boruta). The algorithm ran for 500 iterations (maxRuns) with 500 trees to compute the importance of each taxa. Only selected taxa were retained for final model training and testing.

#### Model development for normal growth maturation

To define a normal growth maturation reference, we trained a Random Forest regression model using the randomForest^48^ package (v4.7-1.2). Infants with consistent normal growth were split into training (80% infants) and testing (remaining 20%) sets to avoid data leakage across longitudinal samples. Model performance was evaluated on the held-out test set using Root Mean Square Error (RMSE) and the R-squared (R^2^). To visualize the contribution of the most important taxa for gut virome or bacteriome maturation, we calculated the correlation (R^2^) between each taxa abundance and infant’s chronological age and plotted these values against Mean Gini Importance. In addition, we generated a heatmap using the pheatmap^49^ package (v1.0.13), displaying the mean abundance of Boruta^47^ selected taxa across age bins, categorized by their species taxonomy and VC identifier. Relative abundance or nucleotide diversity values were scaled by row to highlight temporal shifts in the viral landscape across the first 600 days of life.

#### Maturation model validation and maturity gap analysis across nutritional growth trajectories

The random forest models trained on infants with consistent normal growth were used to predict the microbiome age for all samples across all nutritional trajectories including consistent normal growth (test set), consistent moderate stunting, consistent severe stunting, decline to moderate stunting and recovery to normal growth. To compare microbiome maturation across trajectories relative to the normal growth group, we fitted Generalized Linear Mixed Models (GLMM) using the R package lme4^50^ (v2.0-1) with a Gamma distribution and log-link function to the predicted microbiome age as a function of the infant’s chronological age in the normal growth test group. subjectID was included as a random intercept to account for repeated measurements. This model defined the reference normal growth trajectory. Then, for each sample we calculated the maturity gap, which quantifies the deviation of the microbiome development from the normal growth maturation reference, defined as follows:

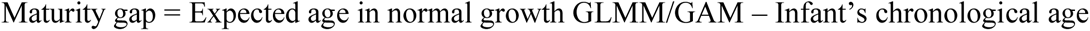

For statistical comparison, the maturity gap of malnourished and transitioning groups was compared against the healthy reference using Wilcoxon rank-sum tests. Negative gap values were interpreted as gut microbiome immaturity.

#### Data visualization and statistical analysis

Data visualization was primarily performed using the ggplot2^51^ package in R (v4.0.2), while maturation heatmaps were generated using the pheatmap^49^ package (v1.0.13). Statistical comparisons between groups were conducted using pairwise Wilcoxon rank-sum tests, with p-values adjusted for multiple testing using the Bonferroni correction. Significant differences (p ≤ 0.05) were identified and annotated directly on plots using the stat_compare_means() function from the ggpubr^52^ package (v0.6.3), with non-significant comparisons omitted. Correlation analyses were performed using the non-parametric Kendall’s Tau coefficient. All visualizations employed color-blind–friendly palettes to ensure accessibility.

## Results

### Most infants are moderately stunted and show dynamic growth trajectories

Considering only infants with mothers with a negative HIV-status^14^ (See Methods), we used the LAZ values reported on the day of sampling to assign a growth status to each sample: normal growth, moderate stunting, or severe stunting. We report significant differences in LAZ across all growth categories for both male and female infants (**Figure 1b**; p<0.001, Wilcoxon test). None of the children were acutely malnourished (wasted) and there are no significant differences in WHZ for any nutritional group or sex (**Figure 1b**; p>0.05, Wilcoxon test). Infant mid-upper arm circumference (MUAC) values indicated that some infants in every nutritional group fell below the 125mm threshold established by the World Health Organization (WHO) to identify infants at high risk of mortality and morbidity^53^ (**Supplementary Fig 1A**). This suggests that even infants showing normal growth may have reduced fat and muscle reserves, highlighting an elevated underlying risk of malnutrition in this population.

We then determined the overall growth trajectories for each infant and identified five growth trajectories (**Supplementary Fig 1B;** see Methods). Infants with consistently normal growth included 92 males and 74 females, while infants with consistently moderate stunted growth comprised 7 males and 11 females. Five males and 4 females showed consistently severe stunted growth. The remaining infants showed dynamic growth trajectories that included infants who recovered to normal growth (6 males only), and infants whose trajectory declined from normal to moderate stunted growth (12 males and 8 females) (**Supplementary Fig 1**).

### Rare and transient vOTUs dominate the infant gut virome and reflect interactions with early life gut bacteria

Of the 22,937 vOTUs recovered, 263 vOTUs were classified as complete (1.15%), 1,488 as high quality (6.5%), and 1,411 as medium quality (6.15%) using CheckV^30^. Within the completely assembled vOTUs, 114 vOTUs were identified as virulent (43.3%) and 149 temperate using BACPHLIP^36^. In the same way, 2,039 vOTUs were identified as temperate (70.3%) the high and medium quality vOTUs. The remaining 860 vOTUs were predicted as virulent because they did not contain the canonical genes required for lysogeny. As this could be due to incomplete contig assembly, we classified these vOTUs as incomplete non-temperate (**Figure 2a**). We then predicted the vOTU host families^37,38^ for 91.5% of all vOTUs (n=21,043), which represented 117 distinct bacterial families. Notably, only 7 bacterial families accounted for 75% of all host-assigned contigs, with *Lachnospiraceae* and *Bacterioidaceae* alone representing 40% of host-assigned contigs, indicating a strong viral association with hallmark early life gut bacterial taxa (**Figure 2b**).

**Figure 2.**
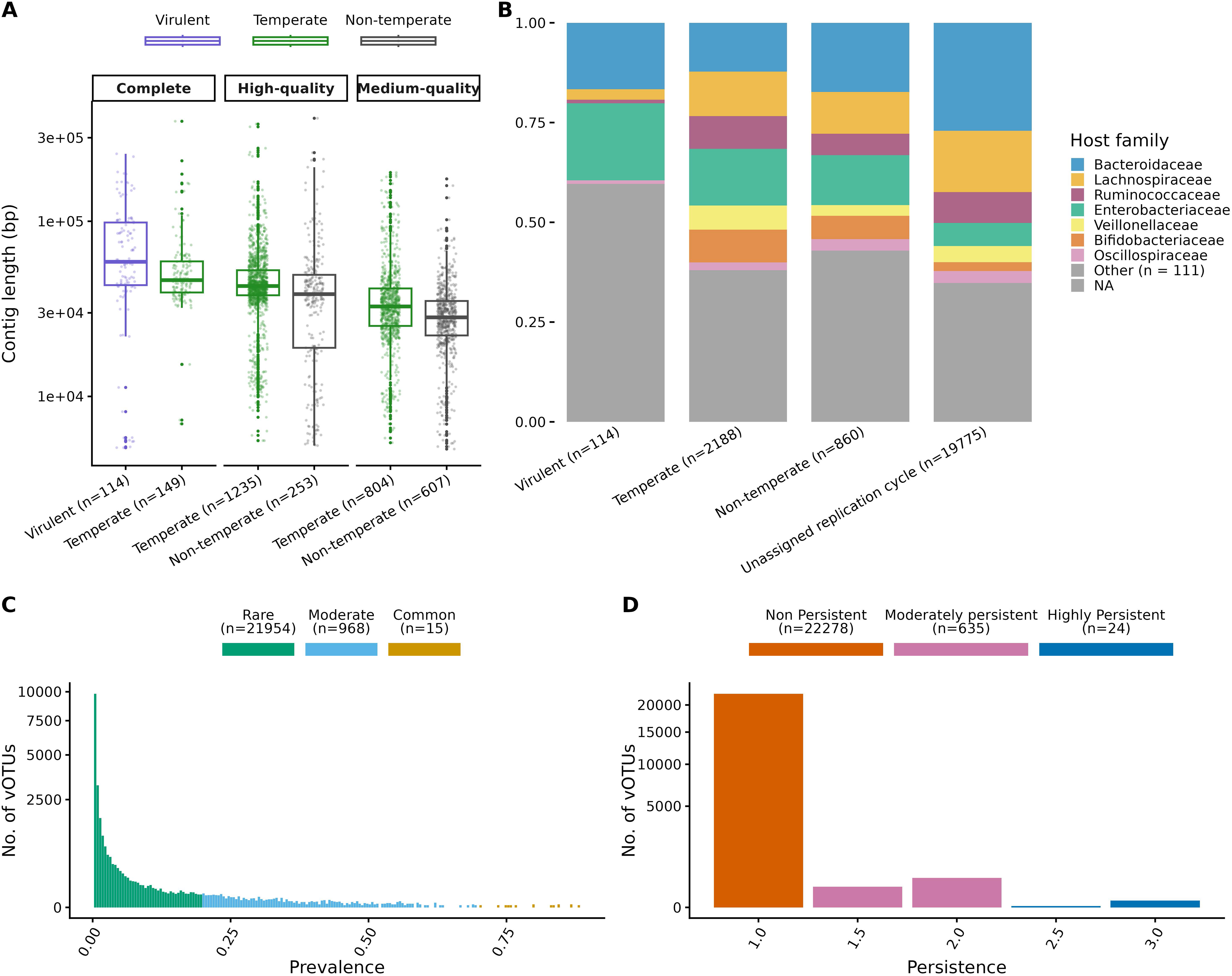
Infant gut virome assembly, quality control, prevalence and persistence. **(A)** Length distribution of the assembled and dereplicated vOTUs across CheckV^30^ quality categories. Of the 22,937 total vOTUs, only those with 50% completeness (n = 3,162) were used for replication cycles prediction via BACPHLIP^36^. vOTUs were classified as virulent only if they were complete, non-complete vOTUs predicted as virulent were categorized as “incomplete non-temperate”. (B) Distribution of predicted bacterial hosts families (stacked bars) for all the 22,937 vOTUs across replication cycle annotations. (C) Histogram of vOTU prevalence based on the percentage of infants in which they were present (Rare: <20%, Moderate: 20%-80%, Common >80%). (D) Histogram of vOTU persistence showing the average number of timepoints a vOTUs was detected within the same infant (Non-persistent: <1, Moderately persistent: 1-2, Highly persistent >2.5).

Prevalence analysis of these vOTUs showed that the viral community was primarily composed of rare vOTUs (n = 21,954; 95.7%), defined as vOTUs detected in <20% of infants. Moderately prevalent vOTUs, defined as detected in 20-70% of infants, accounted for 4.2% of the vOTUs (968 vOTUs); while only 15 vOTUs appeared in >70% of infants and can be considered common (**Figure 2c**). Within each infant, 97.1% vOTUs were non-persistent (n = 22,278), appearing only at one timepoint. Transient vOTUs were present across 1.5-2.5 timepoints and corresponded to 2.8% total vOTUs (n=635), while only 24 vOTUs (0.1%) were considered as persistent as they were detected in more than 3 time points (**Figure 2d**). The average number of days between samples and detection of vOTUs ranged widely, from 10 days to nearly 600 days post-birth, suggesting some local stability despite substantial turnover of viral communities during infancy.

### Bacterial and temperate phage diversity increase with age, regardless of infant growth trajectories

Given the importance of temperate phages for seeding the infant gut virome and their long-term associations with bacterial hosts^11,12^, we focused all subsequent analyses on temperate phages. To describe alpha diversity dynamics in both temperate phage and bacterial communities, we applied Generalized Additive Mixed Models (GAMMs; **Figure 3**). Temperate phage Shannon diversity was modeled as a function of infant age and the model explained 57% of the variance in the dataset. In contrast, the bacterial Shannon diversity GAMM model explained 74% of the data’s variance. Together, these results support the individualistic nature of the virome.

**Figure 3.**
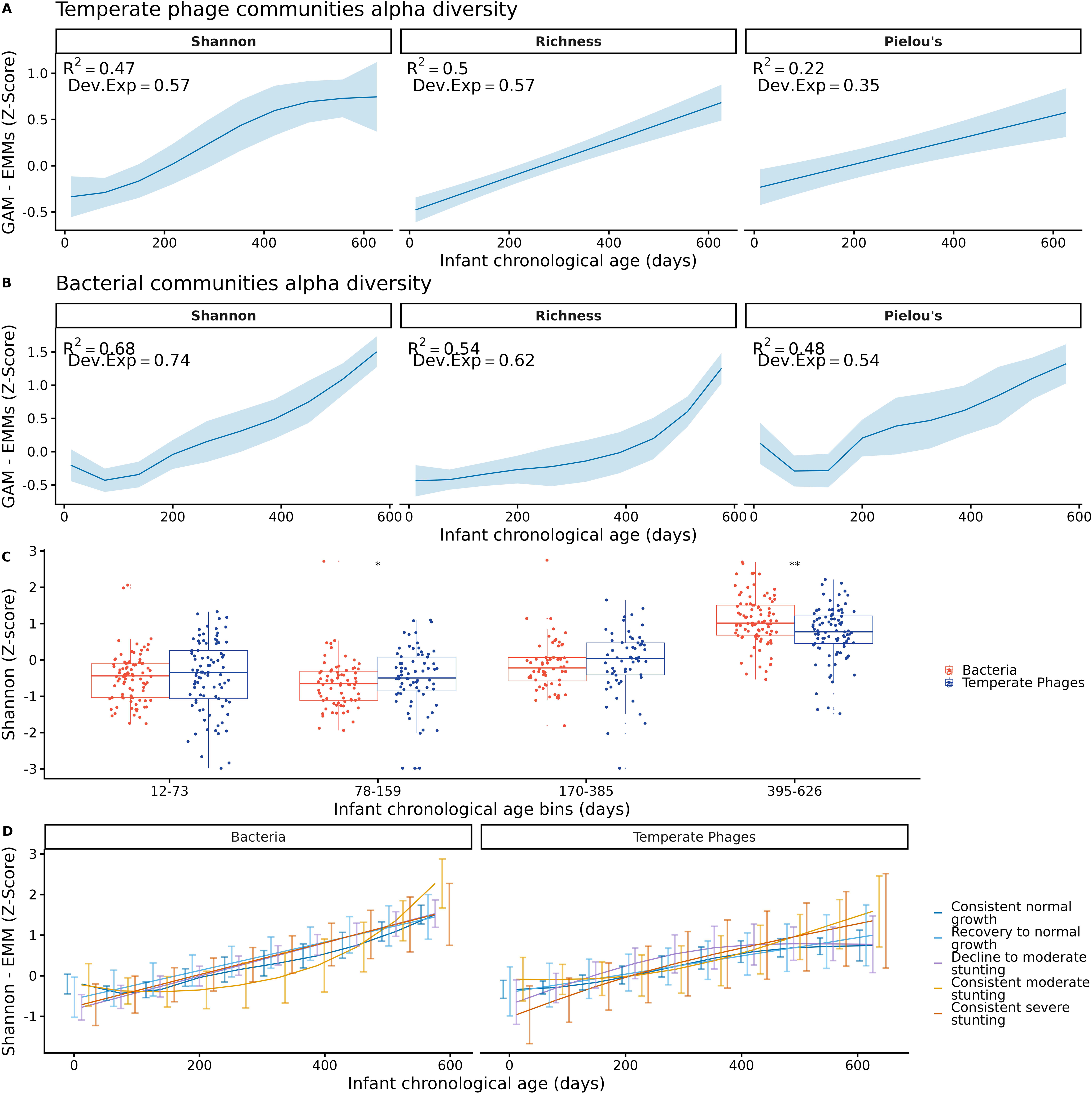
Alpha diversity trajectories of the infant gut virome and bacteriome. **(A)** GAMM modeling of Shannon diversity (AIC = 1,042.1) and its components: richness (observed vOTUs; AIC = 1,000.6) and evenness (Pielou’s index; AIC = 1200.7) within the temperate phage of infants with normal growth. (B) Corresponding GAMM models for Shannon diversity (AIC = 754.4), richness (observed bacterial species; AIC = 953.2), and evenness (AIC = 991.2) within bacterial communities of infants with normal growth. (C) Z-scaled Shannon diversity values for bacterial and temperate phage communities. Groups were compared across four age bins (defined by k-means clsutering) using Wilcoxon rank-sum tests (∗p<0.05,∗∗p<0.01,∗∗∗p<0.001). (D) Estimated Marginal Means (EMMs) of Shannon diversity trajectories for bacterial and temperate phage communities across distinct infant growth trajectories.

Further analyses on alpha diversity patterns revealed distinct temporal dynamics in temperate phage and bacterial communities. For temperate phages, both the richness (number of vOTUs) and evenness of vOTUs (Pielou’s index) increased with age, indicating a progressive expansion of the temperate phage community. Similarly, Shannon diversity increases with age and reaches a plateau around 400 days, suggesting the community starts to stabilize around this time (**Figure 3a**).

For bacterial communities, both richness and evenness also increased over time, consistent with previous findings^54,55^. Unlike the temperate phage community, bacterial Shannon diversity displayed a transient decrease in early days (before day 200), likely driven by the dominance of few bacterial taxa before the establishment of new ones. This is supported by the rapid increase in bacterial richness observed after day 400, suggesting a rapid influx of new bacterial taxa with more diversified environmental sources and exposures (**Figure 3b**).

Scaled Shannon values for bacteria and temperate phages were compared across age bins for infants with a consistently normal growth trajectory. Significant differences between the Shannon diversity of these two communities were detected in the 78–159 day age bin, consistent with a relatively slow influx of bacterial taxa during early infancy compared with the more rapid accumulation of temperate phages. The difference in Shannon diversity of these two communities was also significant in the 395–626 day bin, aligning with the divergence inferred from the GAMMs, where temperate phage Shannon diversity reached a plateau while bacterial Shannon diversity continued to increase (**Figure 3c**). Across the five growth trajectories, both bacterial and temperate phage communities exhibited increasing Shannon alpha diversity with age. For bacteria, the GAMM smooth terms for age indicated a significant age effect in every infant growth trajectory (p<0.001; **Figure 3d, Supplementary Table 1**). Similarly, for temperate phages, we observe an increase in Shannon alpha diversity in all the growth trajectories (p<0.001; **Figure 3, Supplementary Table 1**).

To directly compare Shannon diversity for both microbial communities, we used our GAMMs to compute Estimated Marginal Means (EMMs) of Shannon diversity along with 95% confidence intervals across age and growth trajectories (See Methods). Overall, Shannon diversity did not differ between growth trajectories (**Figure 3d**; Tukey-adjusted pairwise EMMs at each timepoint). Only infants with consistently normal growth versus those in decline to moderate stunting at 12-80 days post-birth were statistically different (p= 0.017). Therefore, these results support similar alpha diversity patterns despite differences in sample size, growth trajectories, and virome individuality.

### Abundance-based microbiome maturation modeling reveals similar bacterial and viral development across growth trajectories

To further investigate microbial colonization dynamics in the infant gut for bacterial and viral communities, we implemented random forest models (RFMs) to predict infant age based on taxonomic abundance matrices, referring to this prediction as the microbiome age^15^. Comparing the gut microbiome age to infant chronological age across growth trajectories enabled us to build bacterial and viral maturation curves, which we modeled using GAMMs. Given that the majority of vOTUs in our cohort are rare (**Figure 2c**), we grouped our vOTUs into viral clusters (VCs) based on shared protein content. This approach also enabled us to perform taxonomic annotation of VCs based on protein content similarity to viral sequences in their reference database.

To assess model performance and establish a healthy gut microbiome maturation reference to compare across growth trajectories, we trained RFMs using 80% of infants from the normal growth trajectory and validated on the remaining 20% (See Methods). The bacterial model showed a high fit (R^2^= 0.759, **Figure 4a**), whereas the viral model exhibited a modest fit (R^2^=0.496; **Figure 4a**).

**Figure 4.**
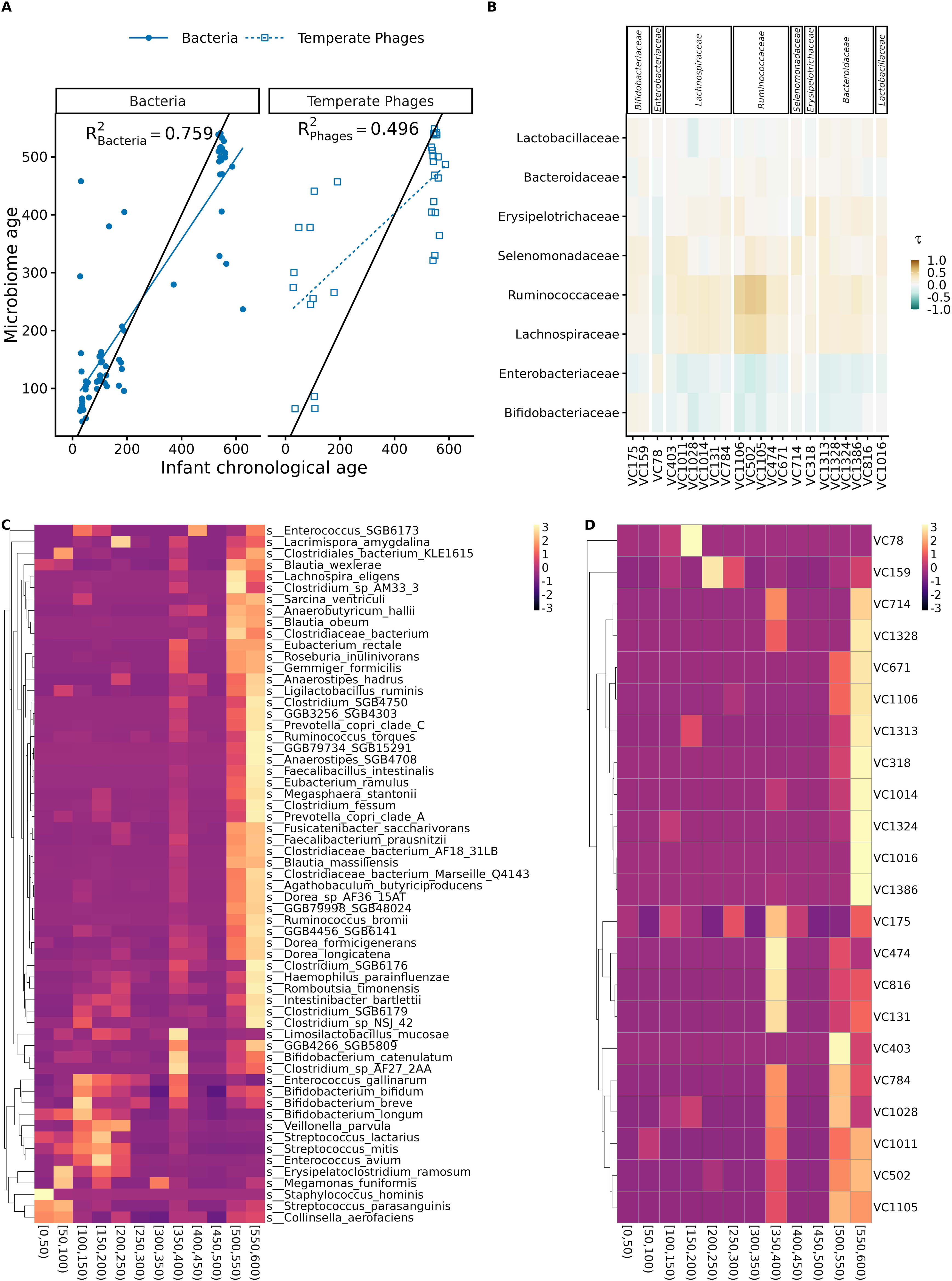
Abundance-based maturation of the infant gut virome and bacteriome. (A) Random forest predicted age based on community composition vs chronological age for both bacterial and temperate phage communities. R^2^ scores indicate the predictive accuracy and fit for each model. (B) Longitudinal Kendal’s Tau correlation coefficients between the average relative abundances of bacterial families and temperate phage viral clusters (VCs) over time in infants with normal growth. VCs are grouped by the family of their predicted bacterial host. Positive tau values indicate co-occurrence, while negative values represent inverse abundance patterns. (C) Heatmap of Z-scaled average relative abundances for age-discriminant bacterial species over time in infants with normal growth. (D) Heatmap of Z-scaled average relative abundances for age-discriminant temperate phage VCs over time in infants with normal growth.

Correlation analyses using Kendall’s Tau revealed coupling between temperate phages and their bacterial hosts across infancy (**Figure 4b**). Overall, VCs exhibited positive correlations with the abundance of their predicted bacterial hosts. The different levels of correlation indicate heterogeneous phage–host relationships that vary by bacterial family. Notably, VCs targeting *Bifidobacteriaceae* displayed negative correlations with early colonizers like *Enterobacteriaceae,* but had a positive correlation with bacteria prevalent beyond the weaning period. Our RFMs allowed us to identify the taxa most predictive of age, and by assessing their correlation with age over time, we identified early and late colonizers within each community. Among bacteria, early settlers included *Bifidobacterium longum* and *Staphylococcus hominis,* while *Faecalibacterium prausnitzii*, *Blautia wexlerae* appeared later. Notably, we detected a species replacement among *Bifidobacterium* species: the abundance of the early colonizer *Bifidobacterium longum* decreased with age as it was replaced by *B. bifidum*, *B. breve,* and *B. catenulatum*, each showing a positive correlation with age (**Figure 4c, Supplementary Fig 2a**).

Consistent with these correlations, patterns of Z-normalized mean relative abundance across age bins revealed temporal shifts in both communities (**Supplementary Fig 3**). VCs associated with early colonizing bacterial families such as *Bifidobacteriaceae* and *Enterobacteriaceae* were most abundant during early infancy. In contrast, VCs linked to late-colonizers, including *Lachnospiraceae* and *Ruminococcaceae,* increased predominantly at later time points, coinciding with the weaning-dependent maturation of microbial communities. Together, these results indicate that the maturation of temperate phage communities parallels bacterial succession in the infant gut, with transitions from early- to late-stage succession patterns observed across both communities. As VC host predictions are made at the family level^38^, we did not observe the replacement in *Bifidobacterium* species detected in bacterial maturation models. However, all our VCs targeting *Bifidobacteriaceae* had a positive correlation with age (**Figure 4b,d, Supp. Fig 2B**).

We then applied our RFMs trained on infants with consistently normal growth to infants from all the other growth trajectories to determine whether there were differences in microbial maturation. For both bacterial and temperate phage communities, the maturity gap (defined as microbiome age - infant’s chronological age) did not differ significantly across growth trajectories (p>0.05, Wilcoxon test, consistent normal growth as reference group). These results indicate that microbial maturation is comparable across all growth trajectories (**Figure 5a,b**), suggesting that the maturation of bacterial and temperate phage communities are not affected by chronic malnutrition or stunting. This contrasts with the previously reported microbial delay associated with wasting resulting from acute malnutrition^15^.

**Figure 5.**
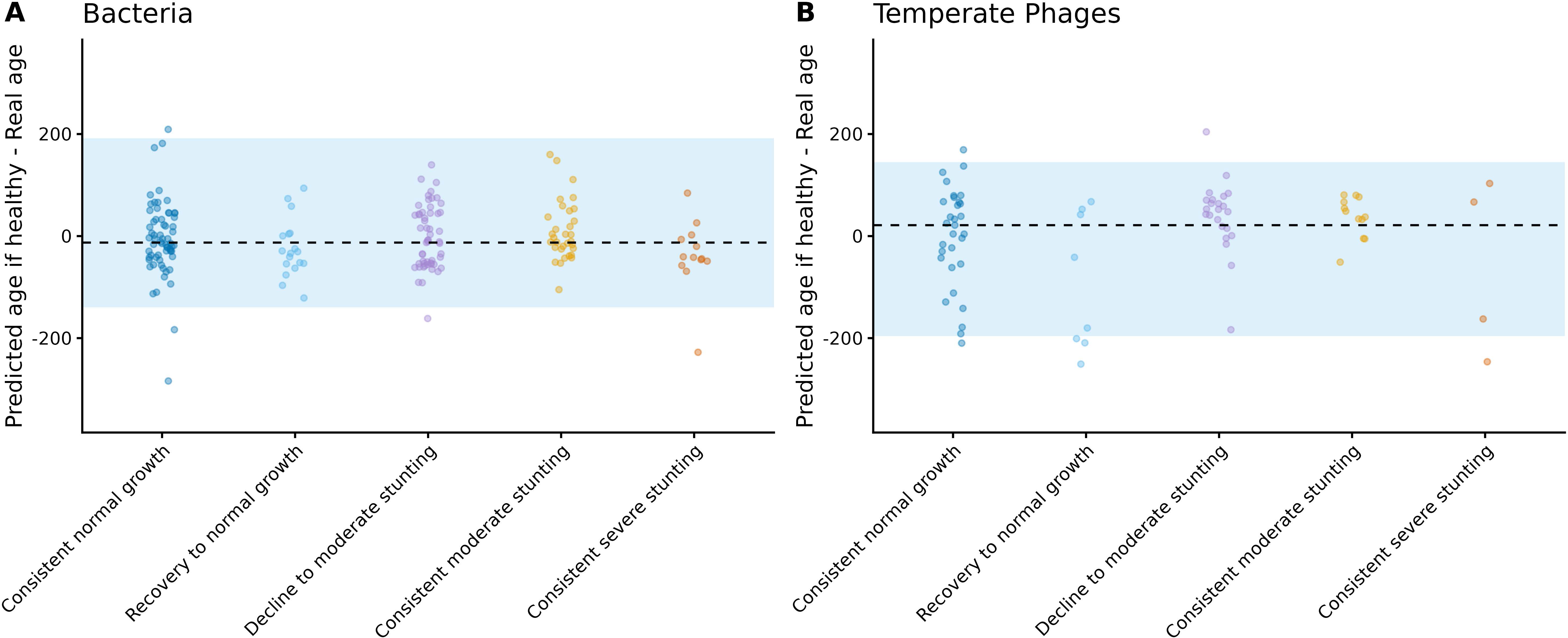
Maturation gap of the infant gut virome and bacteriome across growth trajectories. **(A)** Deviation of the predicted bacteriome age from chronological age using the model trained on the infants with normal growth trajectories. (B Deviation of the predicted temperate virome age from chronological age using the model trained on the infants with normal growth trajectories. In both panels, the blue area represents 95% confidence interval of the maturation gap in infants with normal growth as a reference group. The dashed black line indicates the median gap for the reference group. Positive values indicate accelerated maturation, while negative values indicate a delayed maturation. Maturation gaps were compared across growth trajectories using Wilcoxon rank-sum tests, only significant comparisons (∗p<0.05,∗∗p<0.01,∗∗∗p<0.001) are shown.

Although our abundance-based temperate phage maturation model revealed broadly similar developmental patterns across infant growth trajectories, these VC-level patterns may mask subtle biological differences. Indeed, our model showed a modest fit (R^2^=0.496), suggesting that VC-level abundance alone may not fully capture viral developmental dynamics. These limitations raise the possibility that the effects of stunting could become apparent at finer vOTU strain-level analyses. We thus examined viral strain-level dynamics by tracking nucleotide diversity over time and testing whether these patterns differ between normal growth and stunted trajectories.

### Nucleotide diversity reveals maturation signals masked by abundance-based models

As read-depth variance correlated with age (GLMM, p=0.0816), we first assessed if nucleotide diversity was also dependent on read depth^56^. We report a significant correlation, but with an effect size close to 0 for all nutritional statuses, and we therefore did not consider its effect (**Supplementary Fig 4**). Because all vOTUs in the top predictive VCs had broad nucleotide diversity distributions, we trained the nucleotide diversity model using vOTUs as predictors (**Figure 6a**). Virome maturation modeled according to vOTU nucleotide diversity showed improved age prediction performance (R^2^=0.683, **Figure 6b**) than our abundance-based model.

**Figure 6.**
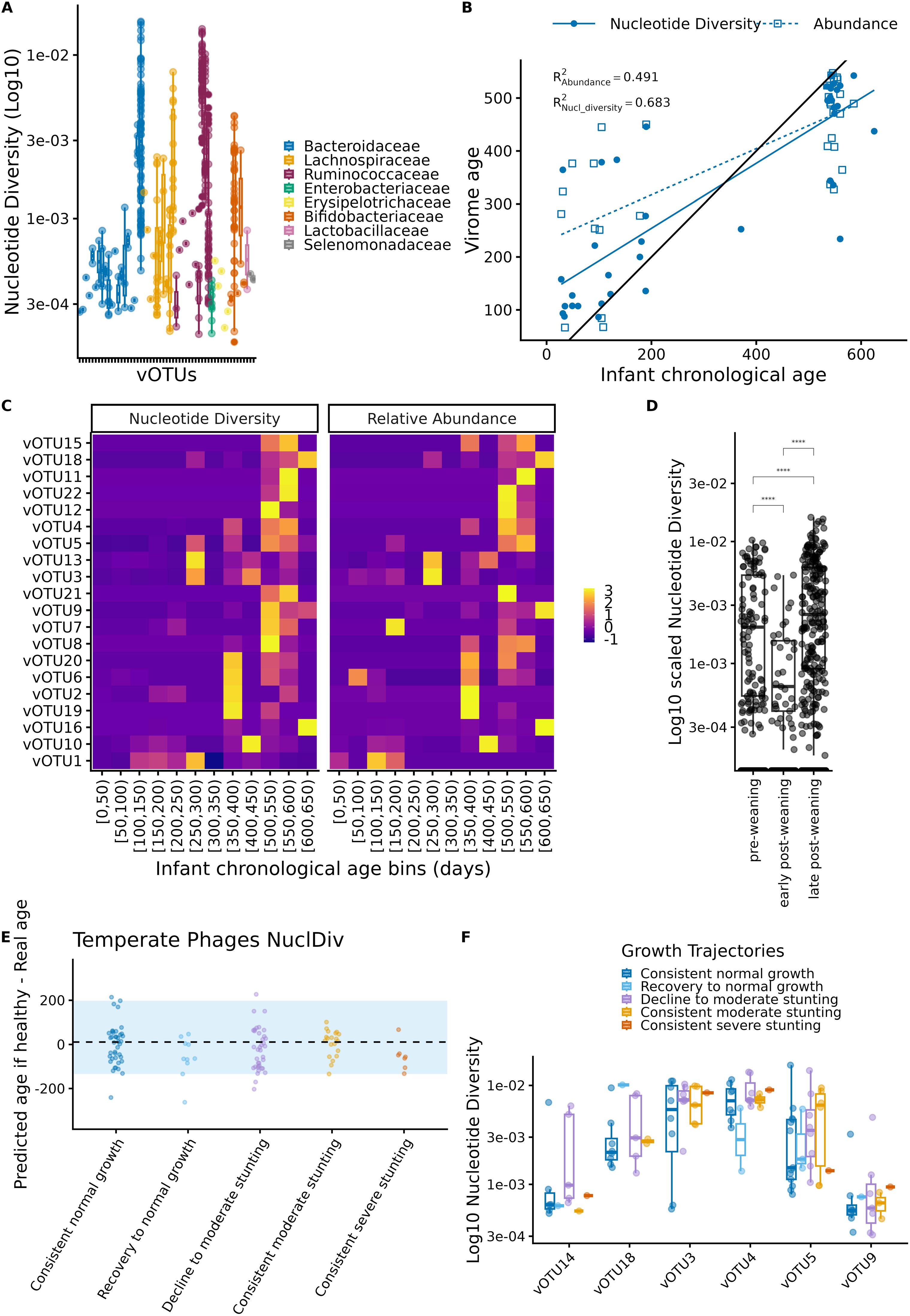
Nucleotide diversity-based maturation of the infant gut virome. **(A)** Log_10_ scaled nucleotide diversity distribution across vOTUs comprising age-discriminant temperate VCs, color-coded by their predicted bacterial host family. (B) Random forest predicted age based on either virome composition (relative abundance) or microdiversity (nucleotide diversity) vs chronological age. R^2^ scores indicate the predictive accuracy and fit for each model. (C) Dual heatmap of Z-scaled average nucleotide diversity (left) and relative abundance (right) for age-discriminant vOTUs over time in infants with normal growth. Rows are matched between heatmaps to illustrate the relationship between virome composition and genetic diversity. (D) Log_10_ scaled nucleotide diversity distribution across pre-weaning, early and late post-weaning periods. (E) Deviation of the predicted virome age from chronological age using the nucleotide diversity-based model trained on the infants with normal growth trajectories. The blue area represents 95% confidence interval of the maturation gap in infants with normal growth as a reference group. The dashed black line indicates the median gap for the reference group. Positive values indicate accelerated maturation, while negative values indicate a delayed maturation. (F) Log_10_ scaled nucleotide diversity distribution across the distinct growth trajectories for age-discriminatory vOTUs. For all distribution plots, metrics in the y-axis were compared across categories in x-axis using Wilcoxon rank-sum tests, only significant comparisons (∗p<0.05,∗∗p<0.01,∗∗∗p<0.001) are shown.

When assessing the most predictive vOTUs, nucleotide diversity increased gradually across chronological age bins, with most vOTUs displaying low nucleotide diversity in early infancy followed by marked increases around weaning (∼200days, **Figure 6c**). In contrast, vOTU relative abundance showed different dynamics, indicating that changes in nucleotide diversity were not necessarily accompanied by corresponding changes in abundance for a given vOTU (**Figure 6c**). To quantify their relationship, we used Kendall’s Tau to correlate vOTUs nucleotide diversity and abundance, identifying that changes in microdiversity are not solely driven by shifts in vOTU abundance and could capture an independent signal of virome maturation (**Supplementary Fig 5**). Notably, we detected correlations between nucleotide diversity of one vOTU and the abundance of other vOTUs, revealing patterns that suggest phage-phage interactions spanning across both compositional (ecological) and genomic-variation (evolutionary) dimensions of the temperate phage community (**Supplementary Fig 5**).

At the community level, nucleotide diversity increased with age (**Supplementary Fig 6;** GAMM - p=7.48E-13) and exhibited distinct maturation phases corresponding to pre-weaning, early post-weaning, and late post-weaning periods (**Figure 6d**). Consistent with our previous findings, we detected significant pairwise differences in nucleotide diversity across weaning phases (Wilcoxon tests; p < 0.05; **Figure 6d**). Nucleotide diversity was lowest during the pre-weaning and early post-weaning phases, and reached its highest levels in the late post-weaning period. Furthermore, phage vOTU diversity (macrodiversity) and microdiversity matrices were significantly correlated (Mantel test; Bray–Curtis distances, r = 0.428, p =0.001), indicating that communities diverging in phage species diversity also diverged in within-phage genomic variation (**Supplementary Fig 7**).

To assess differences in microdiversity maturation, we examined the maturation gap across all the growth trajectories. We report that nucleotide diversity maturation gaps were similar across growth trajectories (**Figure 6e**). The only exception was for the group of infants recovering to normal growth, as they showed a modest but significant delay in nucleotide diversity maturity compared to infants with consistently normal growth (p=0.0401, Wilcoxon test). However, when focusing on the six vOTUs most predictive of age in the nucleotide diversity model (**Supplementary Fig 8**), no significant differences in nucleotide diversity were observed across growth trajectories (**Figure 6f**), although nucleotide diversity distributions varied between growth trajectories. Together, these results demonstrate that temperate phage nucleotide diversity follows a structured, age-dependent maturation that parallels bacterial maturation and is mostly unaffected by growth impairment.

## Discussion

Early life represents a critical window for gut microbial communities as they assemble and mature in parallel with their host. Bacterial succession during infancy has been thoroughly characterized^1,2,15^, but substantially less is known about the phage community and in particular how temperate phage communities fit into this process and whether altered nutritional states modify phage-host interactions^9,11,12^. In this study, we leveraged part of the bulk fecal metagenomes from the SHINE study^23^ to assess how temperate phage communities assembled and evolved during the first 600 days of life^14^. We showed that temperate phages also follow an structured, age-dependent maturation tightly linked to early-life bacterial colonizers, and that their viral microdiversity captures maturation signals masked when using only abundance-based models. We further report that within this cohort, stunting resulting from chronic undernutrition was not associated with altered bacterial or viral maturation trajectories.

Temperate phage communities predominantly target bacterial families associated with early life, highlighting a close coupling between phages and their hosts during gut microbial colonization^38^. Although infant viromes were highly individualized, with most vOTUs being rare and non-persistent, a subset of temperate phages persisted across the entire sampling period (∼600 days), indicating that some temperate phages can be maintained despite extensive turnover in both bacterial and viral community composition^57^. Specifically, these persistent phages may be retained within persistent early bacterial hosts, such as *Bifidobacteriaceae,* and contribute to virome stability despite the bacterial species replacement we observed for *Bifidobacterium* species. Such dynamics parallel findings from stable microbial ecosystems, where species-level composition is conserved while bacterial genomic variability accumulates in accessory functions related to phage defense and resource utilization^58^.

Both bacterial and temperate phage alpha diversity increase with age, consistent with a progressive establishment of microorganisms during infancy but contrasting with what has been reported for virome alpha diversity^13,38^. These studies have focused on virus-like particles (VLPs), predominantly capturing extracellular phages that contract due to low initial bacterial biomass^59^. In contrast, our analysis targets temperate prophages, whose diversity is coupled to bacterial colonization and expansion, and therefore increases as bacterial communities establish. Consistent with this interpretation, a recent infant virome meta-analysis identified significant increases in viral richness during the first months of life, suggesting that early virome diversification may be more pronounced when accounting for methodological, geographical and temporal differences across studies^60^. However, bacterial and temperate phage maturation trajectories are not entirely concordant, suggesting that the viral community development is not a mere reflection of bacterial abundance. The weaning period emerged as a critical transition point for microbial maturation dynamics, coincident with shifts in nutrient availability, turnover from early milk-associated colonizers to taxa adapted to complex glycans from solid dietary substrates, and changes in phage diversity, highlighting how bottom-up forces have a direct effect on bacterial and temperate phage communities^20^.

A central finding of this study is that viral genomic microdiversity is a previously underexplored dimension of early-life virome development. Nucleotide diversity increased with age across many temperate phages and provided stronger maturation signals than abundance-based models, indicating that virome development is also driven by within-phage population diversification. This is similar to what has been observed in bacterial communities, where strain-level variation has been associated with key early life phenotypes, such as birth mode and milk utilization capacity^43,61^. This increase in nucleotide diversity over time is consistent with the idea of diversity-begets-diversity^62^, posing that expanding host and phage species-level diversity creates ecological opportunities that promote within-phage genomic diversification through niche partitioning and host-phage interactions^19,62^. However, this effect was vOTU-specific, suggesting heterogenous eco-evolutionary strategies among members of the temperate phage community. Given that small genomic variation can alter phage host range or trigger host jumps^63–65^, this suggests that viral microdiversity may play an active role in shaping bacterial composition during early microbiome assembly, an example of a top-up effect^20,58^.

Contrary to the delayed maturation of bacterial and viral communities reported in wasting or acute malnutrition^9,15^, we did not observe any alteration or delay in microbial maturation in stunting, irrespective of the models assessed (i.e., abundance- or microdiversity-based). This underscores important biological differences between the different forms of undernutrition and their intersectionality^66^. Stunting is an intergenerational multifactorial condition linked to the mother’s and infant’s health, along with socioeconomic factors^67^. These data demonstrate that stunting still occurs in the absence of an impaired microbiome development or maturation, suggesting adequate microbiome-dependent nutrient absorption function for these infants.

Several limitations should be considered in this work. First, the sampling density of the original study may limit sensitivity to transient perturbations, particularly during the weaning period. In addition, infants in this cohort originate from a region with a high prevalence of stunting and long-term undernutrition, which may reduce contrasts in gut microbiome development between nutritional groups^68–70^. Finally, while bulk metagenomes enable high-resolution characterization of prophage dynamics, paired VLP-enriched metagenomic data would be required to better distinguish between active and dormant prophage states, which are hypothesized to play a key role in gut virome development^11,12^. Despite these limitations, our study expands current understanding of early-life gut microbiome assembly by identifying viral genomic microdiversity as an informative and previously underexplored dimension of virome maturation. Together, these findings suggest that infant gut microbial colonization and succession are shaped by the interplay of top-down forces exerted by phages and bottom-up constraints related to bacterial resources and host availability, operating predominantly at the strain level. Incorporating viral microdiversity metrics into future studies of infant gut microbiome development will therefore be essential for capturing early-life microbial dynamics and their relevance for long-term host health.

## Data and code availability

Code used for data analyses is available at https://github.com/lcamelo10/SHINE_temperate_virome_bacteriome_maturation.

## Acknowledgements

We thank the families and infants enrolled in the SHINE cohort for their participation and contribution to this study. This work was funded by a Canadian Institute of Health Research project grant (PJT-175065) awarded to C.F.M. L.C.C.V is supported by the Fonds de Recherche Quebec – Nature et Techcnologies scholarship (FRQNT; #368509) and a Digital Research Alliance of Canada fellowship (DRAC #265495). We are grateful to the Quebec Centre for Biodiversity Science (QCBS) for funding the internship in the BCEM lab under the supervision of A.R.M and for providing travel awards for the presentation of these data.

We also thank the members of the Maurice Lab, BCEM Lab and Shapiro Lab for their support and advice and Dr. Jesse Shapiro for his guidance as a member of my doctoral advisory committee. We also acknowledge the DRAC in Canada and Direccion de Servicios de Informacion y Tecnologia (DSIT) at Universidad de los Andes in Colombia for providing high-performance computing resources essential to this work.

L.C.C.V: Conceptualization, data curation, formal analysis and funding acquisition (Digital Research Alliance of Canada Champion, Quebec Centre for Biodiversity Sciences), investigation, methodology, software, visualization and writing - original draft. C.F.M: conceptualization, funding acquisition (CIHR grant), supervision, visualization (Figure 1A schematics), and writing - review and editing, A.R.M: resources (internship support), validation, supervision, writing - review and editing.

## Notes

https://github.com/lcamelo10/SHINE_stunting_temperate_virome_bacteriome_maturation

### Competing Interest Statement

The authors have declared no competing interest.

## References

1. Bonham, K. S. et al. Gut-resident microorganisms and their genes are associated with cognition and neuroanatomy in children. Science Advances 9, eadi0497 (2023).

2. Sawhney, S. S. et al. Gut microbiome evolution from infancy to 8 years of age. Nat Med 31, 2004–2015 (2025).

3. Bäckhed, F. et al. Dynamics and Stabilization of the Human Gut Microbiome during the First Year of Life. Cell Host & Microbe 17, 690–703 (2015).

4. Koenig, J. E. et al. Succession of microbial consortia in the developing infant gut microbiome. Proceedings of the National Academy of Sciences 108, 4578–4585 (2011).

5. De Bruyn, F., James, K., Cottenet, G., Dominick, M. & Katja, J. Combining Bifidobacterium longum subsp. infantis and human milk oligosaccharides synergistically increases short chain fatty acid production ex vivo. Commun Biol 7, 943 (2024).

6. Bui, T. N. Y., Paul, A., Guleria, S., O’Sullivan, J. M. & Toldi, G. Short-chain fatty acids—a key link between the gut microbiome and T-lymphocytes in neonates? Pediatr Res 1–9 (2025) doi:10.1038/s41390-025-04075-0.

7. Fahur Bottino, G., et al. Early life microbial succession in the gut follows common patterns in humans across the globe. Nat Commun 16, 660 (2025).

8. Khan Mirzaei, M., et al. Bacteriophages Isolated from Stunted Children Can Regulate Gut Bacterial Communities in an Age-Specific Manner. Cell Host & Microbe 27, 199–212.e5 (2020).

9. Reyes, A. et al. Gut DNA viromes of Malawian twins discordant for severe acute malnutrition. Proceedings of the National Academy of Sciences 112, 11941–11946 (2015).

10. Zhao, G. et al. Intestinal virome changes precede autoimmunity in type I diabetes-susceptible children. Proceedings of the National Academy of Sciences 114, E6166–E6175 (2017).

11. Garmaeva, S. et al. Transmission and dynamics of mother-infant gut viruses during pregnancy and early life. Nat Commun 15, 1945 (2024).

12. Redgwell, T. A. et al. Prophages in the infant gut are pervasively induced and may modulate the functionality of their hosts. npj Biofilms Microbiomes 11, 46 (2025).

13. Gregory, A. C. et al. The Gut Virome Database Reveals Age-Dependent Patterns of Virome Diversity in the Human Gut. Cell Host & Microbe 28, 724–740.e8 (2020).

14. Robertson, R. C. et al. The gut microbiome and early-life growth in a population with high prevalence of stunting. Nat Commun 14, 654 (2023).

15. Subramanian, S. et al. Persistent gut microbiota immaturity in malnourished Bangladeshi children. Nature 510, 417–421 (2014).

16. UNICEF/WHO/World Bank Group. Levels and Trends in Child Malnutrtion. Joint Child Malnutrition Estimates (JME) 2025. https://data.unicef.org/resources/jme/ (2025).

17. Mulyani, A. T., Khairinisa, M. A., Khatib, A. & Chaerunisaa, A. Y. Understanding Stunting: Impact, Causes, and Strategy to Accelerate Stunting Reduction—A Narrative Review. Nutrients 17, 1493 (2025).

18. Mirzaei, M. K. & Maurice, C. F. Ménage à trois in the human gut: interactions between host, bacteria and phages. Nat Rev Microbiol 15, 397–408 (2017).

19. Gregory, A. C. et al. Marine DNA Viral Macro- and Microdiversity from Pole to Pole. Cell 177, 1109–1123.e14 (2019).

20. Ndiaye, M., Bonilla-Rosso, G., Mazel, F. & Engel, P. Phage diversity mirrors bacterial strain diversity in the honey bee gut microbiota. Nat Commun 16, 9738 (2025).

21. Castledine, M. & Buckling, A. Critically evaluating the relative importance of phage in shaping microbial community composition. Trends in Microbiology 32, 957–969 (2024).

22. Silveira, C. B. & Rohwer, F. L. Piggyback-the-Winner in host-associated microbial communities. npj Biofilms Microbiomes 2, 16010 (2016).

23. The Sanitation Hygiene Infant Nutrition Efficacy (SHINE) Trial Team et al. The Sanitation Hygiene Infant Nutrition Efficacy (SHINE) Trial: Rationale, Design, and Methods. Clin Infect Dis 61, S685–S702 (2015).

24. Bolger, A. M., Lohse, M. & Usadel, B. Trimmomatic: a flexible trimmer for Illumina sequence data. Bioinformatics 30, 2114–2120 (2014).

25. Langmead, B. & Salzberg, S. L. Fast gapped-read alignment with Bowtie 2. Nat Methods 9, 357–359 (2012).

26. Blanco-Míguez, A. et al. Extending and improving metagenomic taxonomic profiling with uncharacterized species using MetaPhlAn 4. Nat Biotechnol 41, 1633–1644 (2023).

27. Nurk, S., Meleshko, D., Korobeynikov, A. & Pevzner, P. A. metaSPAdes: a new versatile metagenomic assembler. Genome Research 27, 824–834 (2017).

28. Camargo, A. P. et al. Identification of mobile genetic elements with geNomad. Nat Biotechnol 42, 1303–1312 (2024).

29. Camacho, C. et al. BLAST+: architecture and applications. BMC Bioinformatics 10, 421 (2009).

30. Nayfach, S. et al. CheckV assesses the quality and completeness of metagenome-assembled viral genomes. Nat Biotechnol 39, 578–585 (2021).

31. Bolduc, B. et al. iVirus 2.0: Cyberinfrastructure-supported tools and data to power DNA virus ecology. ISME COMMUN. 1, 77 (2021).

32. Roux, S. et al. Minimum Information about an Uncultivated Virus Genome (MIUViG). Nat Biotechnol 37, 29–37 (2019).

33. Danecek, P. et al. Twelve years of SAMtools and BCFtools. Gigascience 10, giab008 (2021).

34. Roux, S., Emerson, J. B., Eloe-Fadrosh, E. A. & Sullivan, M. B. Benchmarking viromics: an in silico evaluation of metagenome-enabled estimates of viral community composition and diversity. PeerJ 5, e3817 (2017).

35. R Core Team. R: A language and environment for statistical computing. https://www.r-project.org/ (2024).

36. Hockenberry, A. J. & Wilke, C. O. BACPHLIP: predicting bacteriophage lifestyle from conserved protein domains. PeerJ 9, e11396 (2021).

37. Roux, S. et al. iPHoP: An integrated machine learning framework to maximize host prediction for metagenome-derived viruses of archaea and bacteria. PLOS Biology 21, e3002083 (2023).

38. Shamash, M., Sinha, A. & Maurice, C. F. Improving gut virome comparisons using predicted phage host information. mSystems 10, e01364–24.

39. Bolduc, B. et al. Machine learning enables scalable and systematic hierarchical virus taxonomy. Nat Biotechnol 1–10 (2025) doi:10.1038/s41587-025-02946-9.

40. García-Botero, C. & Reyes, A. phu: Phage Utilities - A modular toolkit for viral genomics workflows. Zenodo 10.5281/zenodo.19489078 (2026).

41. Black, E. J. et al. Virus taxonomy: the database of the International Committee on Taxonomy of Viruses. Nucleic Acids Res 54, D776–D789 (2026).

42. R Core Team. R: A language and environment for statistical computing. https://www.r-project.org/ (2025).

43. Olm, M. R. et al. inStrain profiles population microdiversity from metagenomic data and sensitively detects shared microbial strains. Nat Biotechnol 39, 727–736 (2021).

44. Oksanen, J. et al. vegan: Community Ecology Package. (2026).

45. Wood, S. mgcv: Mixed GAM Computation Vehicle with Automatic Smoothness Estimation. (2025).

46. Lenth, R. V., et al. emmeans: Estimated Marginal Means, aka Least-Squares Means. (2026).

47. Kursa, M. B. & Rudnicki, W. R. Boruta: Wrapper Algorithm for All Relevant Feature Selection. (2025).

48. original), L. B. (Fortran, original), A. C. (Fortran, port), A. L. (R & port), M. W. (R. randomForest: Breiman and Cutlers Random Forests for Classification and Regression. (2024).

49. Kolde, R. pheatmap: Pretty Heatmaps. (2025).

50. Bates, D., et al. lme4: Linear Mixed-Effects Models using ‘Eigen’ and S4. (2026).

51. Wickham, H., et al. ggplot2: Create Elegant Data Visualisations Using the Grammar of Graphics. (2026).

52. Kassambara, A., Business, L. E. (Faculty of E. and, Debrecen, U. of & Hungary). ggpubr: ‘ggplot2’ Based Publication Ready Plots. (2026).

53. Rana, R., Barthorp, H., McGrath, M., Kerac, M. & Myatt, M. Mid-Upper Arm Circumference Tapes and Measurement Discrepancies: Time to Standardize Product Specifications and Reporting. Global Health: Science and Practice 9, 1011–1014 (2021).

54. Younge, N. E. et al. Disrupted Maturation of the Microbiota and Metabolome among Extremely Preterm Infants with Postnatal Growth Failure. Sci Rep 9, 8167 (2019).

55. Amenyogbe, N. et al. Bacterial and Fungal Gut Community Dynamics Over the First 5 Years of Life in Predominantly Rural Communities in Ghana. Front. Microbiol. 12, (2021).

56. Bustos-Caparros, E. et al. Uneven sequencing (coverage) depth can bias microbial intraspecies diversity estimates and how to account for it. ISME Commun 5, ycaf228 (2025).

57. Lou, Y. C. et al. Infant gut DNA bacteriophage strain persistence during the first 3 years of life. Cell Host & Microbe 32, 35–47.e6 (2024).

58. Somerville, V. et al. Strain-level and phenotypic stability contrasts with plasmid and phage variability in water kefir communities. 2025.02.27.640646 Preprint at 10.1101/2025.02.27.640646 (2025).

59. Lim, E. S. et al. Early life dynamics of the human gut virome and bacterial microbiome in infants. Nat Med 21, 1228–1234 (2015).

60. Shamash, M. & Maurice, C. F. The infant gut virome assembles rapidly and predictably over the first three years of life. 2026.02.17.705938 Preprint at 10.64898/2026.02.17.705938 (2026).

61. Gough, E. K. et al. Bifidobacterium longum and microbiome maturation modify a nutrient intervention for stunting in Zimbabwean infants. eBioMedicine 108, 105362 (2024).

62. Madi, N., Vos, M., Murall, C. L., Legendre, P. & Shapiro, B. J. Does diversity beget diversity in microbiomes? eLife 9, e58999 (2020).

63. De Sordi, L., Khanna, V. & Debarbieux, L. The Gut Microbiota Facilitates Drifts in the Genetic Diversity and Infectivity of Bacterial Viruses. Cell Host & Microbe 22, 801–808.e3 (2017).

64. De Sordi, L., Lourenço, M. & Debarbieux, L. “I will survive”: A tale of bacteriophage-bacteria coevolution in the gut. Gut Microbes 10, 92–99 (2019).

65. Hussain, F. A. et al. Rapid evolutionary turnover of mobile genetic elements drives bacterial resistance to phages. Science 374, 488–492 (2021).

66. Thurstans, S. et al. The relationship between wasting and stunting in young children: A systematic review. Maternal & Child Nutrition 18, e13246 (2022).

67. Gough, E. K. et al. Maternal fecal microbiome predicts gestational age, birth weight and neonatal growth in rural Zimbabwe. eBioMedicine 68, (2021).

68. Gou, W. et al. Early-life exposure to the Great Chinese Famine and gut microbiome disruption across adulthood for type 2 diabetes: three population-based cohort studies. BMC Med 21, 414 (2023).

69. Huda, M. N. et al. The impact of early-life exposures on growth and adult gut microbiome composition is dependent on genetic strain and parent-of-origin. Microbiome 13, 143 (2025).

70. Rothschild, D. et al. Environment dominates over host genetics in shaping human gut microbiota. Nature 555, 210–215 (2018).

