## Supplementary figures and images for "Temperate phage microdiversity reflects infant gut microbiome maturation independent of chronic undernutrition"

### Supplementary Fig 1

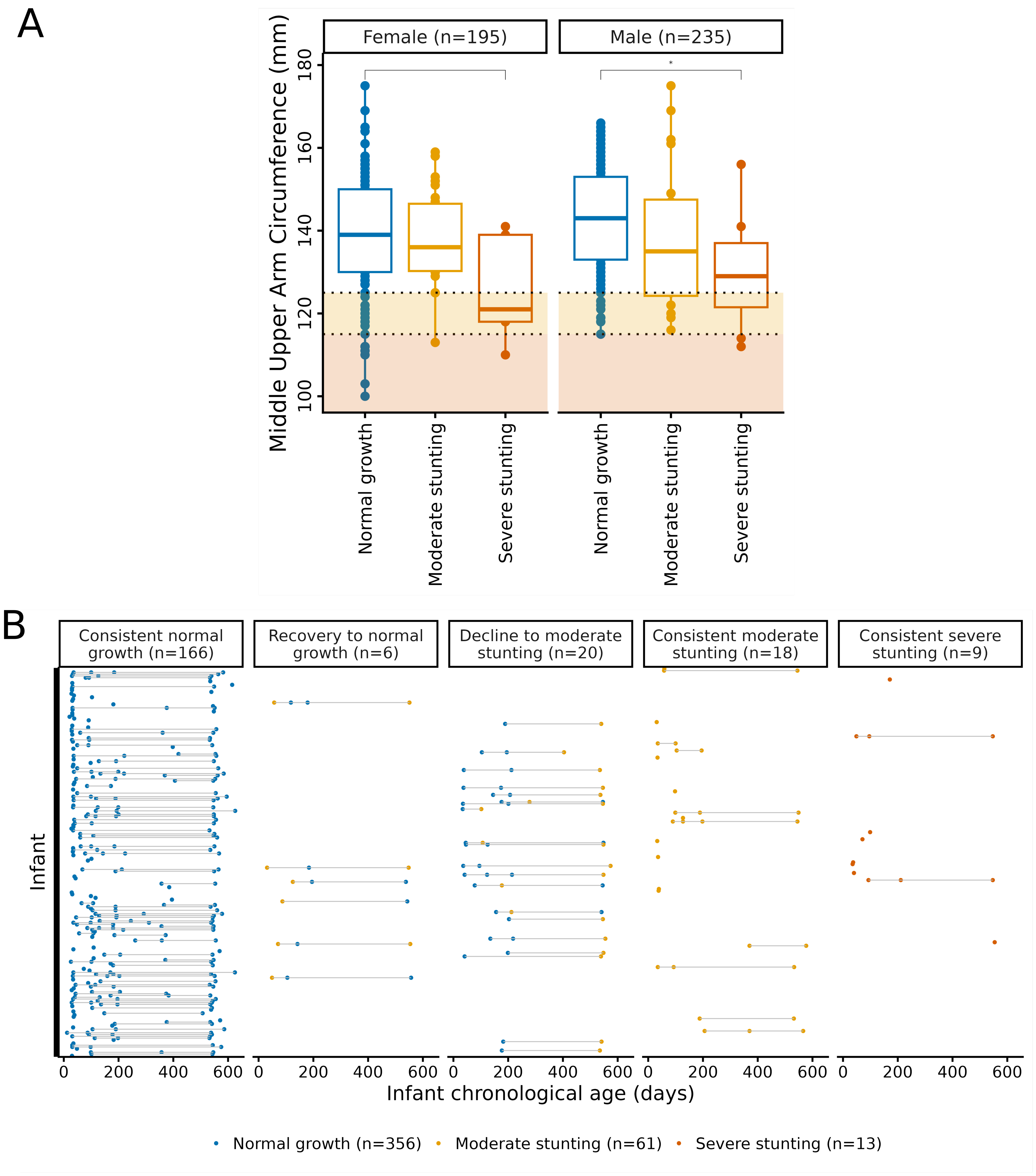

### Supplementary Fig 2

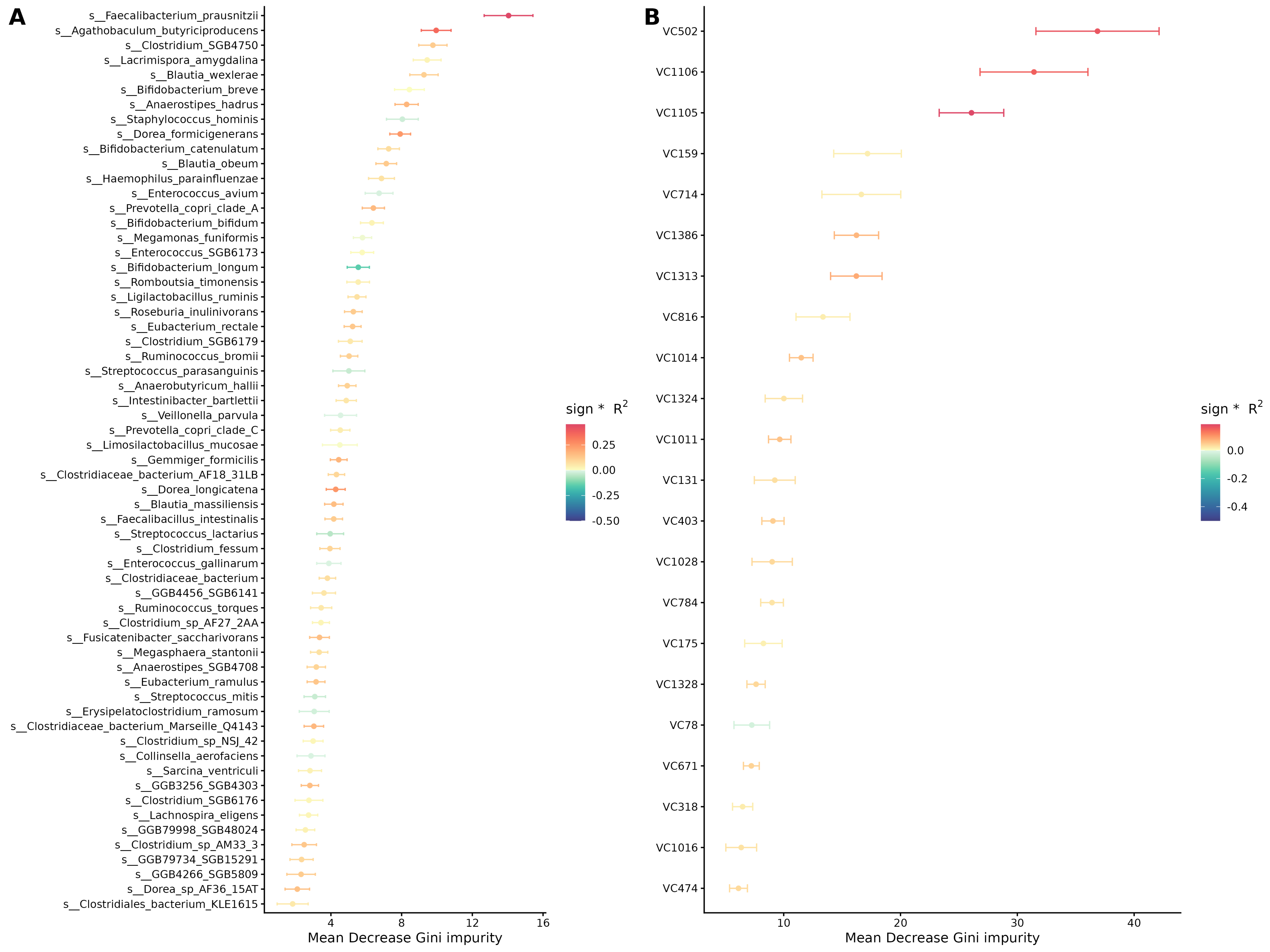

### Supplementary Fig 3

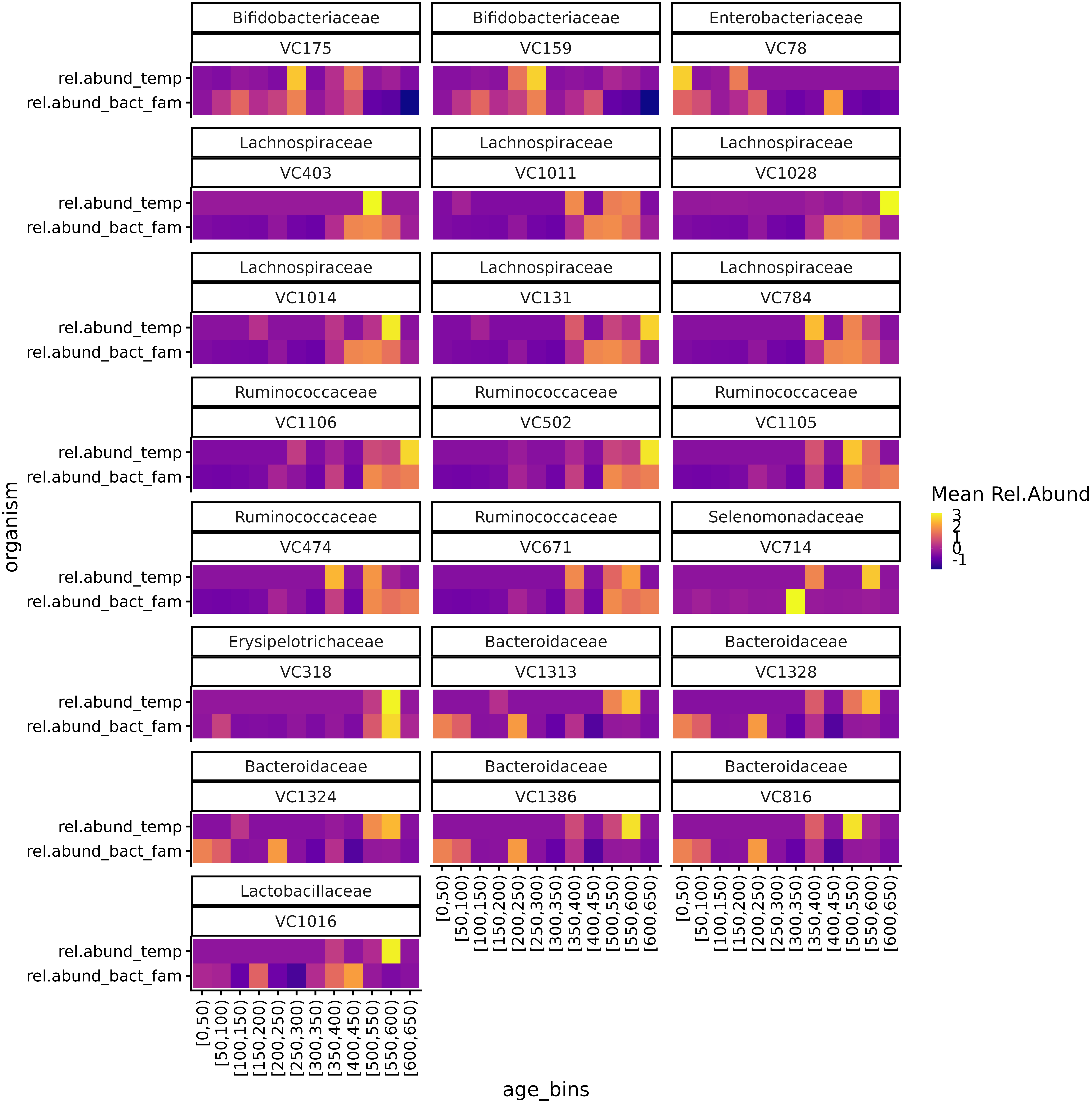

### Supplementary Fig 4

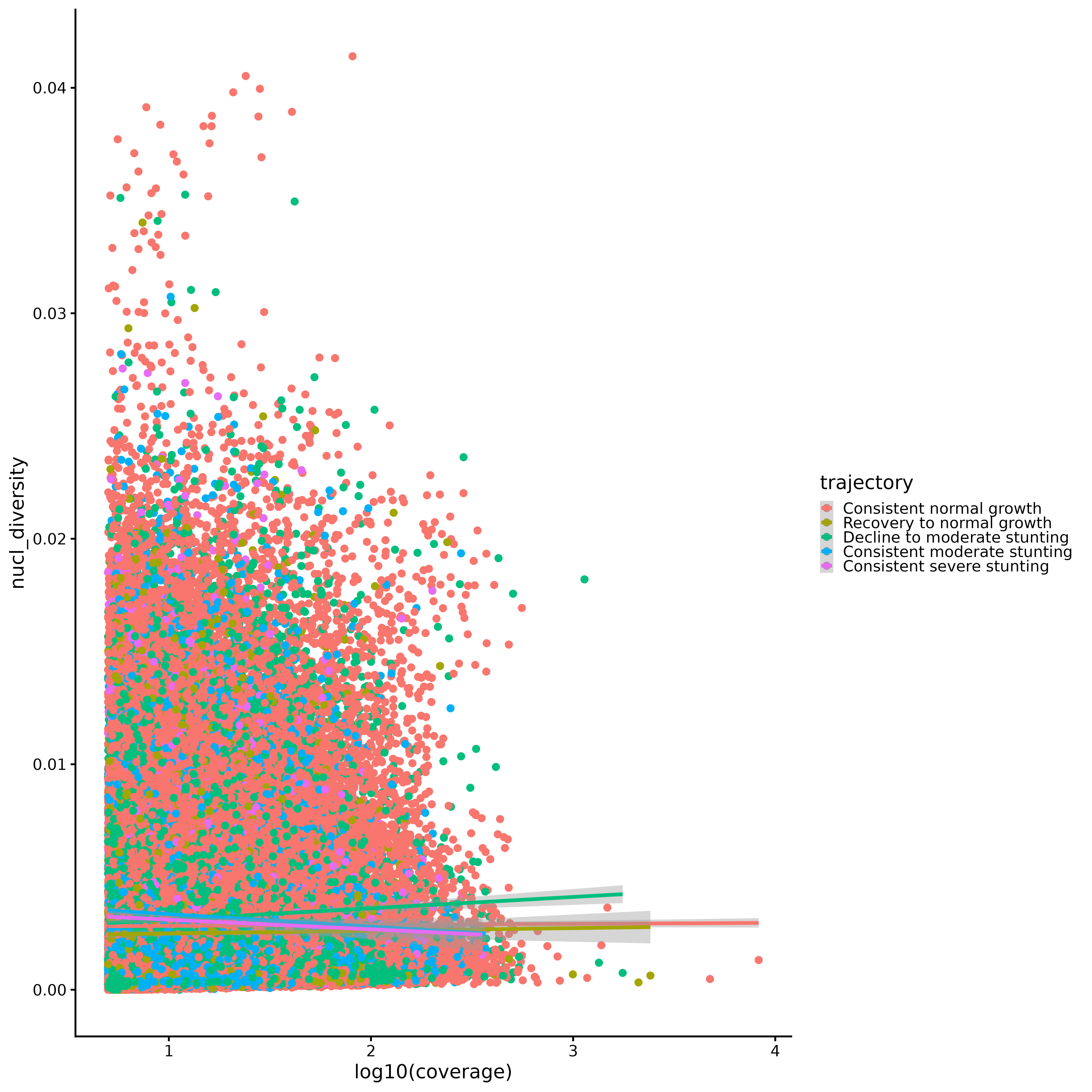

### Supplementary Fig 5

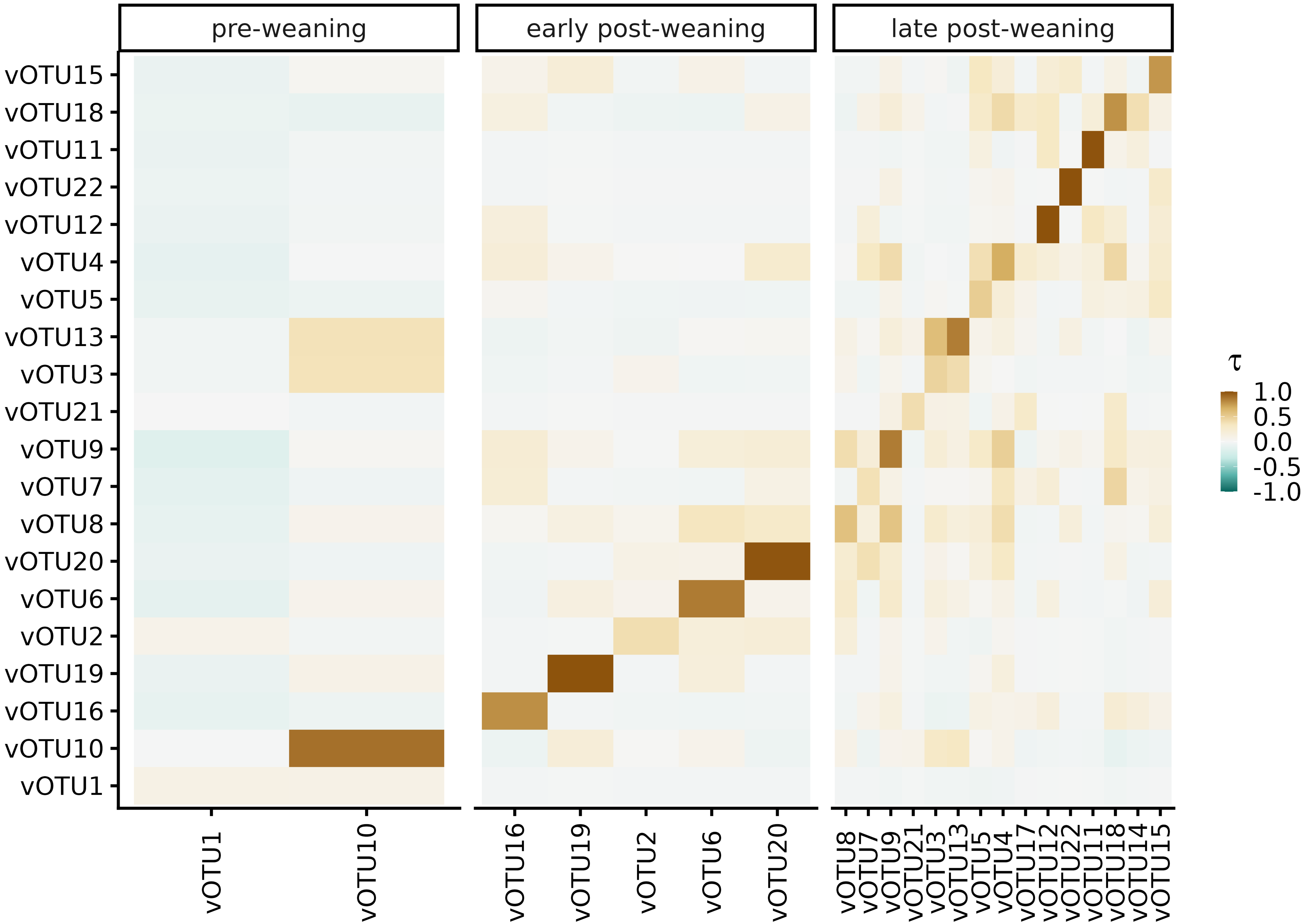

### Supplementary Fig 6

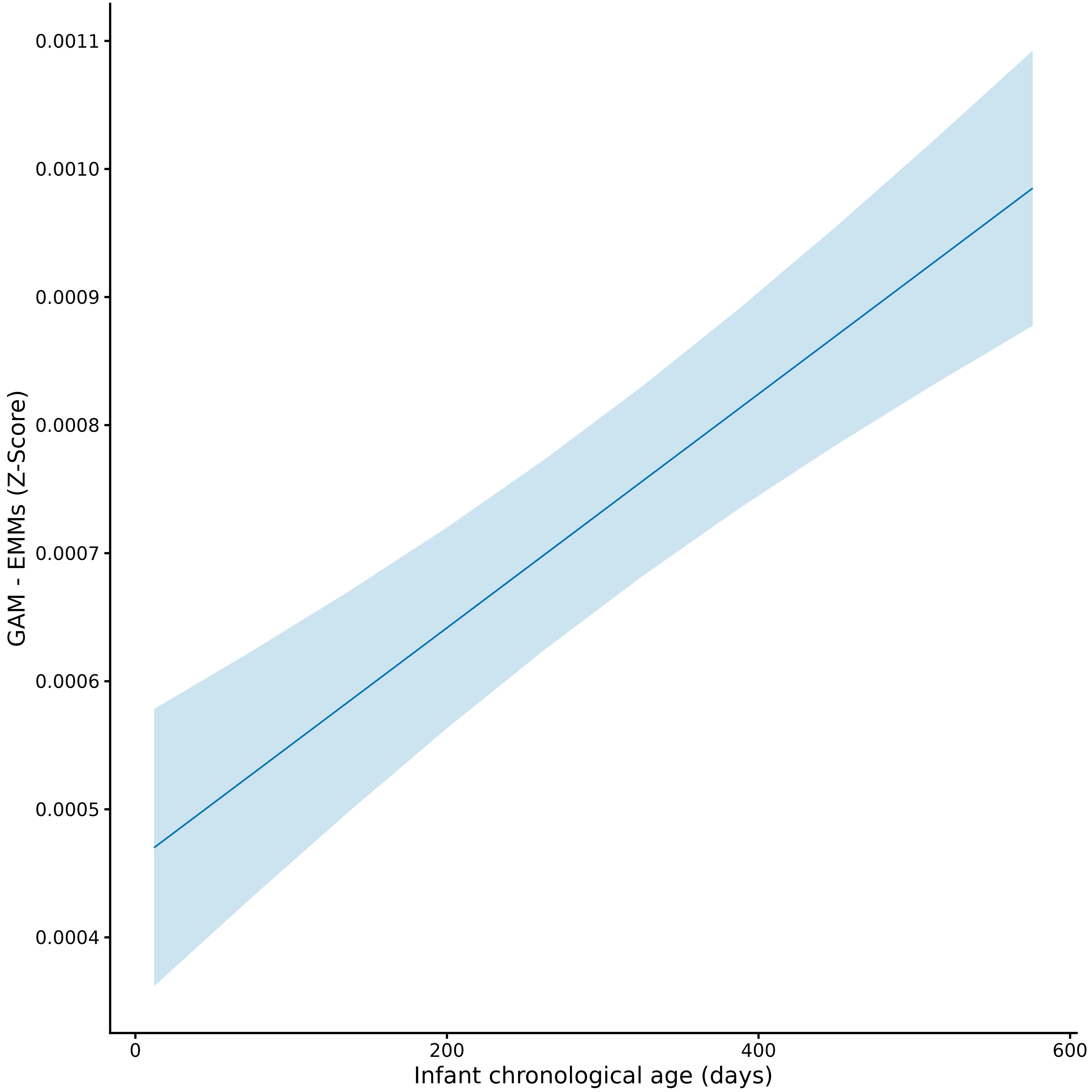

### Supplementary Fig 7

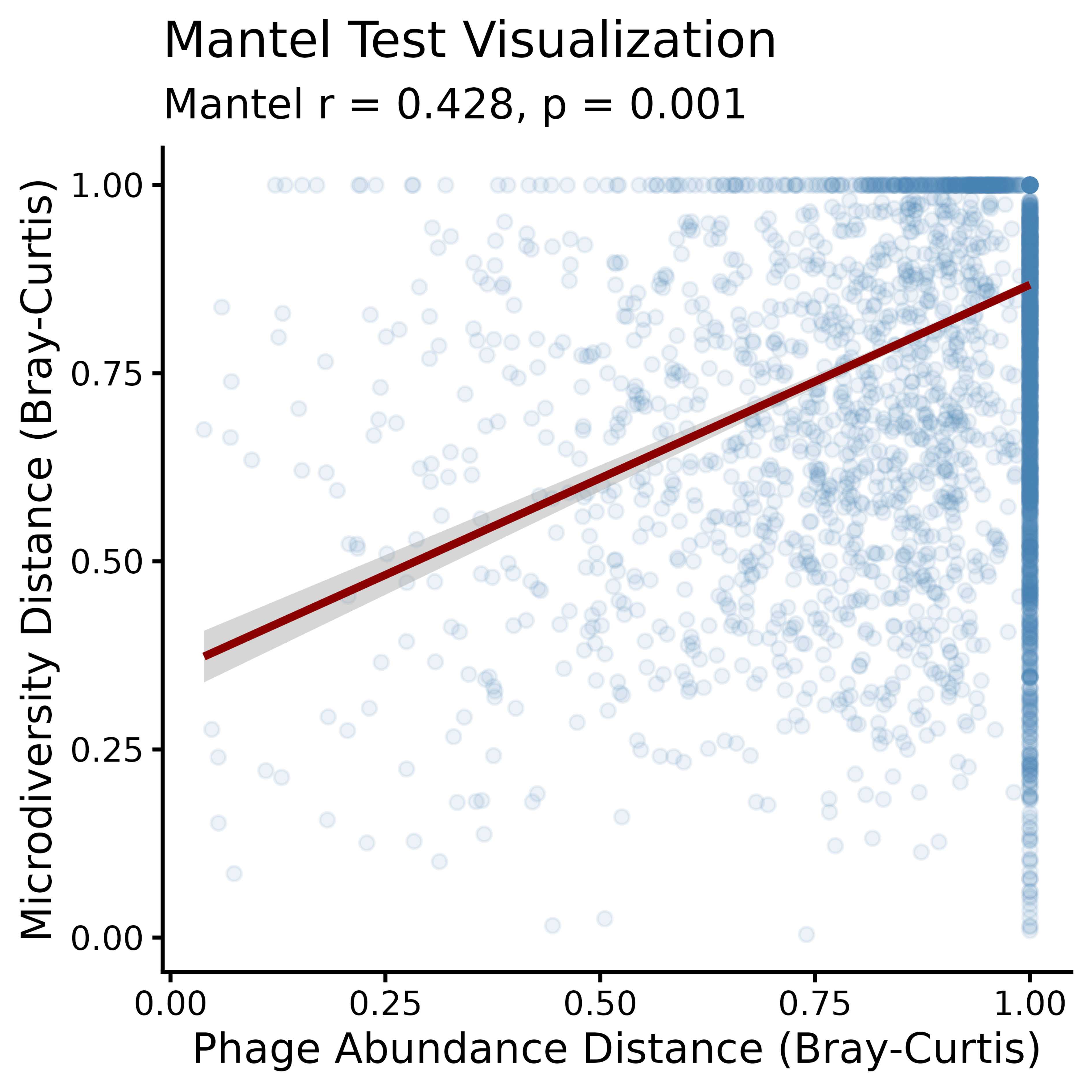

### Supplementary Fig 8

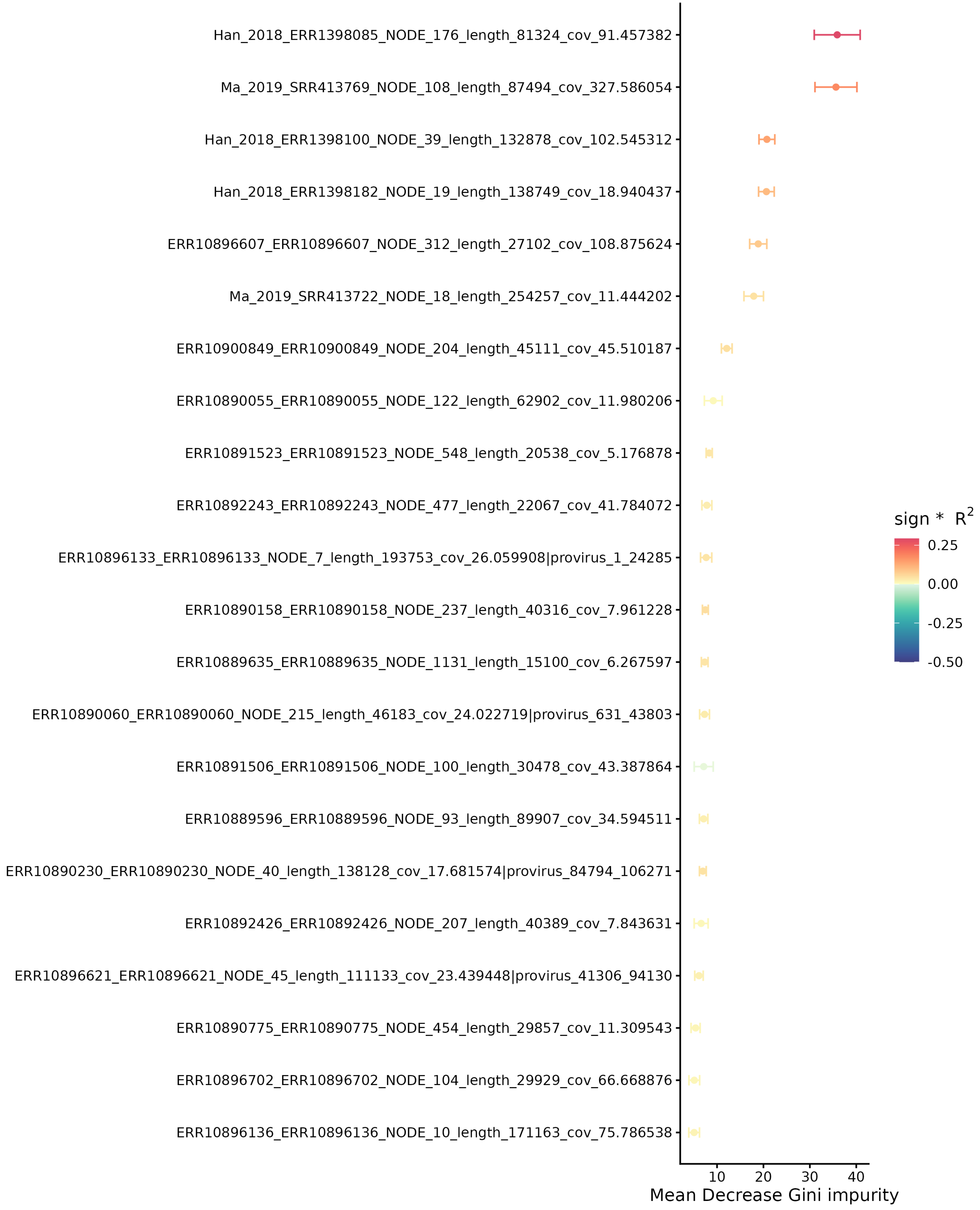
